# Opportunistic partner choice among arctic plants and root-associated fungi is driven by environmental conditions

**DOI:** 10.1101/2024.09.14.613029

**Authors:** B Parisy, N.M Schmidt, A.R Cirtwill, E Villa-Galaviz, M Tiusanen, J Siren, C.F.C Klütsch, P.E Aspholm, K Raundrup, E.J Vesterinen, H Wirta, T Roslin

## Abstract

Interactions between plants and soil microbes play an important role in structuring plant communities. Yet, little is known about how fungal networks are structured on the one hand by fungal responses to their environment (including their host plant) and on the other by responses to each other. We quantified changes in plant–fungus networks along geographic and environmental gradients across the Arctic, assessing the degree to which plants and fungi showed preference for specific interaction partners and how specificity varies along environmental gradients. To this aim, we sampled roots of 12 widely distributed plant taxa: *Saxifraga oppositifolia; Bistorta vivipara; Dryas spp.; Vaccinium tis-idaea; Vaccinium uliginosum; Vaccinium myrtillus; Empetrum nigrum; Betula nana; alix arctica; Salix polaris; Cassiope tetragona;* and *Silene acaulis*. To quantify the pool of fungi from which plant roots may recruit association partners, we also sampled fungi in the surrounding soil. Identifying fungaI communities by DNA metabarcoding, we used Hierarchical Modelling of Species Communities (HMSC) to assess how fungal communities change along environmental gradients, and whether plants actively select their root-associated fungi from the pool of fungi present in the bulk soil. We found that although the fungal communities within the soil and rhizosphere share 85% of genera, their composition differs significantly from each other. The two community types show similar responses to the environment and taxa show low partner fidelity. Thus, the structure of fungal communities on plant rhizosphere is mainly driven by abiotic rather than biotic conditions. Overall, in comparison with null models, networks of plants and rhizosphere-associated fungi showed a distinctly non-random structure, responding strongly to pH and temperature gradients. Our findings suggest that the dynamics and structure of plant-root associated interactions might be severely altered by abiotic changes in the rapidly changing arctic environment.

**Open Research statement:** Data are privately provided for peer review. The raw sequences for the soil and root samples generated during the current study will be available in the Sequence Read Archive repository, in the BioProject PRJNA1094865 upon acceptance. For review purposes, the code and datasets used for the analyses of this study are temporarily available in Figshare open access repository at https://figshare.com/s/1b074f1751682d3487cf upon acceptance.

## Introduction

Belowground microbiomes can affect the composition of plant communities in many ways. For one, root-associated microbial communities can have a strong influence on plant nutrient acquisition as well as affecting the survival of individual plants by improving defence against pathogens (Bever et al. 2010; Friesen et al. 2011; Li et al. 2020). Recent studies have shown that the distributional ranges of plants can be modulated by the presence of suitable mycorrhizal fungi (McCormick et al. 2018, Bahram and Netherway 2021, Romero et al. 2023). Ultimately, ecological dynamics may change plant community composition and diversity, leading to concordant and predictable changes in soil microbial communities and vice versa (Reynolds et al. 2003; Wardle et al. 2004; Bardgett 2011). However, few studies have analysed how root associated fungal communities change along broad regional gradients in the artic realm.

Changes in plant–fungal associations are likely accentuated in the arctic region as the area is rapidly warming (Post et al. 2019; Rantanen et al. 2022), and since arctic plants allocate a large proportion of biomass belowground which likely increases the resource availability for arctic soil microbial communities (Iversen et al. 2015; Eisenhauer et al. 2017; Bowles, Jackson, and Cavagnaro 2018). Furthermore, interactions among and between communities of different kingdoms may change with environmental stress (Classen et al. 2015; Vandenkoornhuyse et al. 2015). However, the precise mechanisms behind (and the microbes involved in) the dynamics of plant-microbe interactions in response to environmental conditions are still poorly understood (Mohan et al. 2014; Trivedi et al. 2022). As a specific prediction regarding general change towards increasingly challenging conditions, the Stress-Gradient Hypothesis (Bertness and Callaway 1994) posits that plant species will show lower specialization in harsher environments. While arctic ecosystems are sensitive to disturbance, there is little information on how soil microbial communities respond to environmental changes (Frindte et al. 2019; Bardgett and Caruso 2020). Current knowledge of how climate change may alter interactions between plants and their root-associated organisms (from being beneficial to antagonistic or vice versa) is still poorly developed.

In this paper, we aimed to fill critical knowledge gaps regarding the direct and/or the indirect effects of abiotic conditions on plant-microbe interactions. Specifically, we aimed to:

1. Assess how the probability of plant-fungus interaction is modulated by the occurrences of the fungi within the soil (Figure 1: Box 1)
2. Disentangle the extent to which plant associated fungi communities are shaped by the biotic and the climatic environment, respectively (Figure 1: Box 2).
3. Evaluate fungal partner selection among plant species by quantifying deviations from randomness in plant–microbe networks across the Arctic (Figure 1: Box 3)

**Figure 1.**
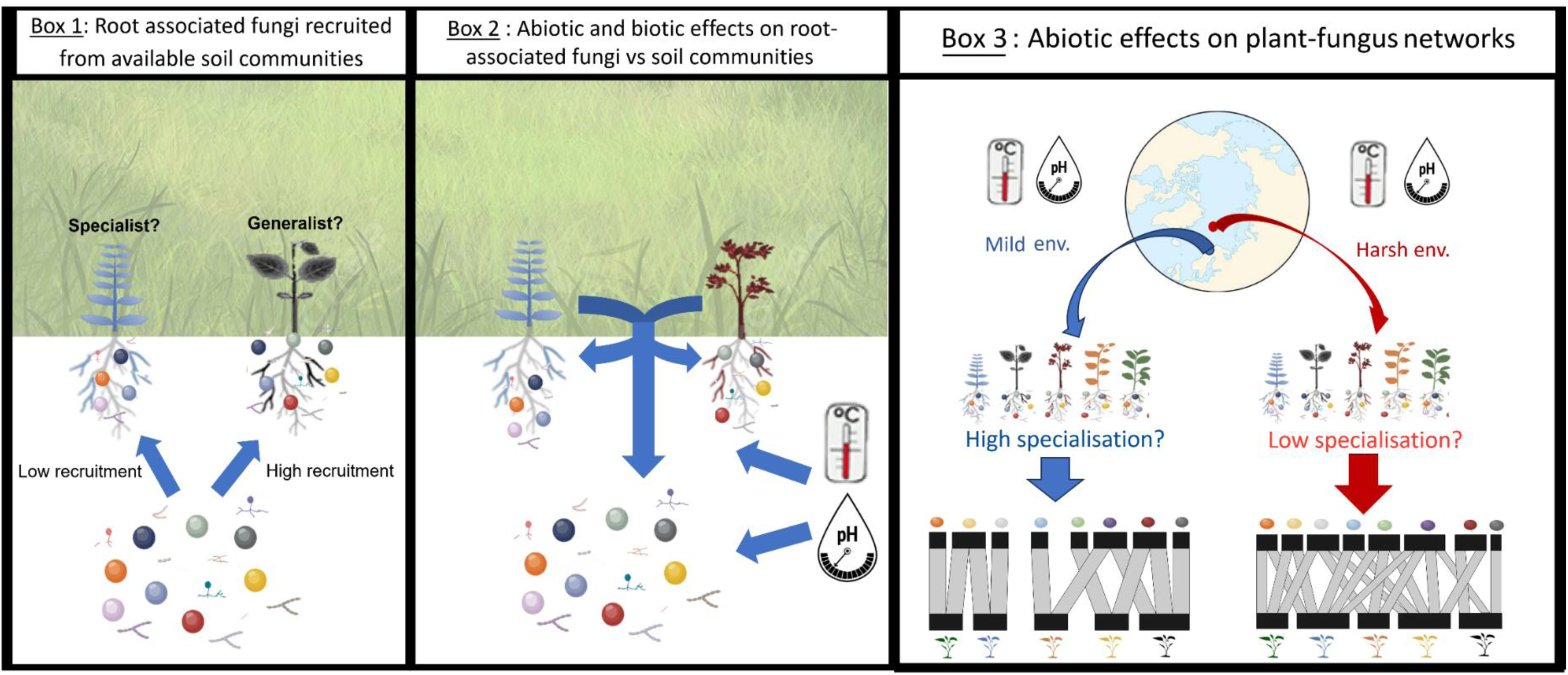
Conceptual setting and approach of this study. To quantify biotic vs abiotic imprints on fungal communities across the Arctic, we examined the relationship between the pool of fungi available in the soil vs the pool of fungi recruited to the rhizosphere (Box 1); the impact of abiotic conditions vs host plant identity on fungi detected in the rhizosphere (Box 2); and changes in partner specificity along geographic and environmental gradients of the Arctic (Box 3).

To this aim, we took a three-fold approach: We first disentangle the effect of the environment from effects of the plants themselves (Figure 1). For that, we compared paired communities to each other: the fungi found associated with the rhizosphere and the fungi found in nearby soil (Figure 1: Box 1). Second, we used a joint species distribution model to quantify the relative importance of the abiotic and biotic environment in shaping the composition of fungi communities (Figure 1: Box 2). Finally, we explored how the associations between plants and fungi change with environmental conditions (Figure 1: Box 3,) by using network analyses applied to plants and their root-associated fungi along environmental gradients.

### *A priori*, we hypothesized the following

a. Based on global patterns (Soudzilovskaia et al. 2015; Bahram et al. 2018; Tedersoo et al. 2022), we expected a strong relationship between the composition of fungi available in the soil and the composition of root-associated fungi. Thus, we expected to observe congruent changes in the composition of these two communities across the Arctic.
b. As an alternative hypothesis to (a), we may expect the plant to provide a pronounced “physical buffer” (Sikes 2010). If so, then we expected to observe differences in how the root associated fungal communities vs. fungi present in the soil respond to environmental gradients. More specifically, we expected that soil fungi will be mainly driven by abiotic conditions, while root-associated fungi will be mainly driven by the identity of the interaction partner.
c. The Stress-Gradient Hypothesis (Bertness and Callaway 1994) predicts increasing ratios of facilitative-to-competitive interactions with increasing stress. Microbes can play a key role in mitigating environmental stress for their plant hosts, and in return, plant can provide physical “shelter” for fungi. From this assumption, we hypothesized that, in harsher environment, where partner availability is likely reduced, the niche breadth of both plant and fungi will be broader. Thus, we expected plant species to show a lower level of network specialization in their microbial associations at high-arctic than at subarctic sites.

## Material and methods

### Sample collection

To evaluate the impact of community assembly processes across multiple spatial scales, we coordinated sampling at two spatial scales: a pan-arctic scale and a regional scale. During the summers of 2020 and 2021, we collected a total of 2900 root and soil samples by sampling seven **sites** across a gradient of 14.5° latitude. These sites ranged over various ecotones from the Subarctic through the Low-Arctic to the High Arctic (Figure 2).

**Figure 2.**
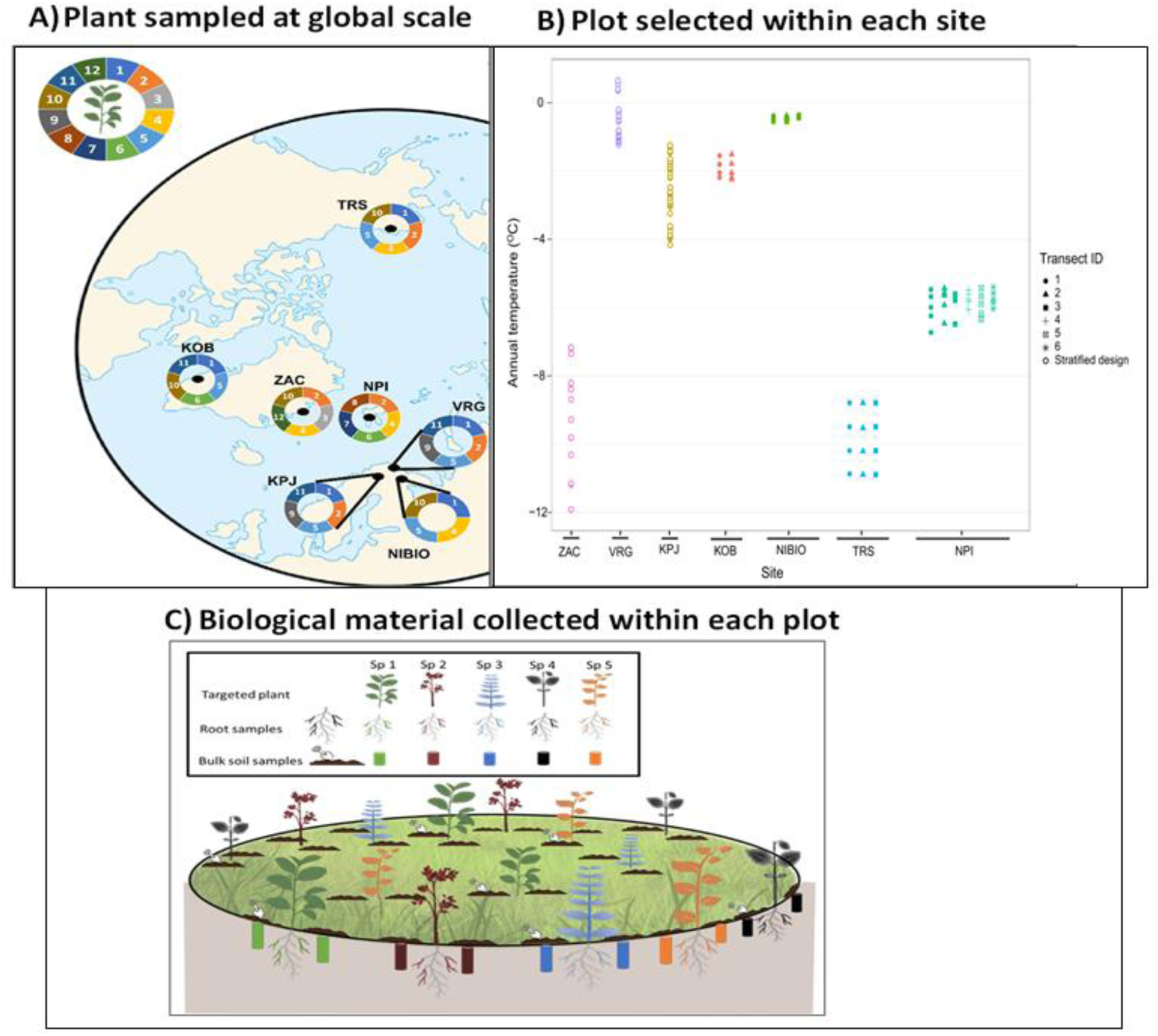
Sampling design and data collection. Panel A) shows our seven sampling sites across the Arctic. At each **site**, we sampled at least 8 **plots** (each with a radius of 25 m) across local elevational gradients. In the graph, each slice of the site-specific doughnut chart represents a focal plant species sampled per site, with the color revealing its identity (see below). Panel B) shows the estimated temperature of the plots selected within each site. The sites are first sorted according to the number of transects locally sampled (from left to right) and then by the number of plots for each site. Panel C) shows the biological sampling design within each plot. We collected roots of three individuals of each target plant as well as two soil cores in the vicinity of each plant. Each plant color represents one of the five plant species targeted per site. Site-specific acronyms: KPJ= Kilpisjärvi, Finland; VRG= Varanger Peninsula, Norway, NIBIO= Gandvik valley, Norway; KOB= Kobbefjord, South-West Greenland; TRS= Toolik Research Station, USA; NPI= Norsk Polarinstitut, Ny Alesund, Svalbard; ZAC= Zackenberg, North-East Greenland. The top-left doughnut chart offers a color legend for the plant species: N°1 = Betula nana; N°2 = Bistorta vivipara; N°3 = Cassiope tetragona; N°4 = Dryas spp (which includes Dryas octopetala, Dryas integrifolia and Dryas octopetala x integrifolia hybrids); N°5 = Empetrum nigrum; N°6 = Silene acaulis; N°7 = Saxifraga oppositifolia; N°8 = Salix polaris; N°9 = Vaccinium myrtillus; N°10 = Vaccinium uliginosum; N°11 = Vaccinium vitis-idaea; N°12 = Salix arctica.

To characterise variation along local environmental gradients within each of our seven sites, we defined at least three local **transects** (Figure 2). These transects were located at least 250 meters apart from each other across a joint elevation gradient. To capture differences in microclimatic environmental conditions, we selected the strongest elevational gradients available in the vicinity of each site (i.e., the gradients spanning the largest difference in metres above sea level). Within three sites (Kilpisjärvi, Varanger and Zackenberg), we sampled more intensively following a stratified random sampling design across multiple elevational gradients (Figure 2).

Along each transect, we established four **plots** with a radius of ca 25 m each. These plots were selected at least 100 metres from each other along the slope. A plot was selected only if at least three on the five target species occurred within the plot (no matter which species; see list below). At each site, we sampled at least 8 plots whereas all geographical coordinates and altitudes were recorded using a GPS (Garmin, GPSmap 62s, Switzerland).

As focal plant species, we selected species common and widespread enough to be sampled across the Arctic region. Thus, within each site, the set of target species to be sampled was defined as the five most locally abundant species out of the following list: *Saxifraga oppositifolia; Bistorta vivipara; Dryas spp; Vaccinium vitis-idaea; Vaccinium uliginosum; Vaccinium myrtillus; Empetrum nigrum; Betula nana; Salix arctica; Salix polaris; Cassiope tetragona; and Silene acaulis* (For the list of species collected within each site, see Figure 2A). Of these taxa, *Dryas* spp. represents a species complex. In North America, the dominant species is *Dryas integrifolia* and in Europe it is *Dryas octopetala*. Nonetheless, the two species interbreed, with individuals in Northeast Greenland (Zackenberg) mainly being hybrids *Dryas octopetala × integrifolia* (Elkington 1965, Philipp and Siegismund 2003).

Importantly, these plant taxa were *a priori* selected to represent mycorrhizal types: *Betula nana, Bistorta vivipara, Dryas spp, Salix polaris and Salix arctica* have been previously assumed to be associated with ectomycorrhiza (ECM; Gardes and Dahlberg 1996, Cripps and Eddington 2005, Abrego et al. 2020); *Cassiope tetragona, Vaccinium myrtillus, Vaccinium uglinosum, Vaccinium vitis-idea and Empetrum nigrum* to be mainly associated with ericoid mycorrhiza (ErM) but sometimes with ectomycorrhiza, too (Treu et al. 1995; Wang and Qiu 2006; Koizumi and Nara 2017; Daghino et al. 2022); *Silene acaulis* and *Saxifraga oppositifolia* to be associated mainly with arbuscular mycorrhiza (AM) but sometimes with ectomycorrhiza (Treu et al. 1995; Wang and Qiu 2006; Fujimura and Egger 2012).

To characterize fungal communities in the rhizosphere (i.e. fungi directly interacting with the target plant species), we collected three roots fragments (length >3 mm) from different parts of the root architecture, for 2-3 individuals of each target plant species found within the plot. To this aim, we gently dug and/or scraped 2-3cm around the focal plant individual until we found the roots (clearly identified as connected to the stem). We then excavated the roots without breaking their fine parts and collected the soil attached to the root. The resulting sample was then considered as a compound community type: the rhizosphere. To characterise the fungal community of the soil, we collected two bulk soil samples on opposite sides of each targeted plant individual (ca 10 cm). For each soil core (ca 5cm diameter), we collected the upper 5-10 cm of the soil layers, with the layers pooled for analysis. Coarse roots (>0.2 mm in diameter) and stones were removed. The soil was then stored within a paper bag and put it into a ZIP-lock bag filled with silica gel, which was then placed at −20°C.

In total, we collected 1450 roots and 1450 soil samples across 129 plots spread across the Arctic (Figure 2).

### Environmental data

To characterise the local climate, we extracted the annual mean temperature (BIO1) of each site from the Worldclim database (Fick and Hijmans 2017). We then associated the average elevation of each plot within a site with a mean annual temperature. Assuming a decline of 0.7°C for every 100m.a.s.l., we used the mean elevation as the baseline and then subtracted or added 0.7°C for every 100m below or above the average, respectively. This 0.7°C factor was defined following the standard lapse rate (Mane et al. 2022) and is consistent with highly-resolved data collected by Peña-Aguilera et al. (2023).

Vegetation cover surrounding each focal plant individual in a 1m² area was estimated visually. The average vegetation coverage of each plot was then calculated from all the individual estimates. For three plots at the Zackenberg site (ZAC10,11 and 12), we lacked information on plot-level vegetation. For these plots, we used the mean of all other plots within the site as a conservative measure of vegetation coverage.

From a part of the soil samples remaining after metabarcoding (see below), we measured soil chemistry as represented by pH, Carbon and Nitrogen content measured at the plot level. From all the soil samples collected within each plot, we pooled soil samples collected around each target plant species sampled locally. The pooled soil was then oven-dried at 70°C and homogenized using a sieve with a mesh size of 2mm. 0.15mg of soil was weighted under air-vacuumed balance, placed in tin foil and analyzed for carbon and nitrogen content using a Leco series 828 series analyzer (Leco, United states). Nitrogen and carbon content was measured for each pooled soil sample using the LCRM method and calibrated with soil samples of known concentration. Another part of the pooled soil sample was used to measure pH following the ISO 10390:2021 standards (International Organization for Standardization; https://standards.iteh.ai/). For this, we prepared a 1:5 (volume fraction) suspension of soil with water, shook the suspension for 60 min using a mechanical shaker and left it to rest at least 1 h before measurement. The pH probe was calibrated with three buffers of pH 4.00, 6.88 and 9.22, respectively.

### DNA metabarcoding

The bulk soil and rhizosphere samples were used for the laboratory analysis, implemented by Bioname Ltd. (www.bioname.fi) as a turn-key service from sample handling through bioinformatics to final data on taxa × sample matrix.

5 ml of the bulk soil samples were transferred to a sterile 50 ml Falcon tube and 1,25 ml of 2,5 mm ceramic beads were added to the tube. The tube containing the sample was homogenized in a Bullet Blender DX50 for 5 minutes. 50–100 mg of the homogenized sample was then used for DNA extraction. The rhizosphere samples were first cut into small pieces using DNA-clean scissors, and then homogenized together with ceramic 2mm beads in sterile 50ml Falcon tube for 10min in a Bullet Blender 50DX (Next Advance, Inc., Troy, NY, USA). DNA was extracted following the protocol of Vesterinen et al. (2016), with some modifications as follows: A fixed volume (30 ml) of pre-warmed 60°C lysis buffer (Aljanabi and Martinez 1997; Vesterinen et al. 2016) and 30 uL of proteinase K were added to the sample, and the mix was incubated for exactly 2 hours 45 min at +60 °C in a shaking incubator. After incubation, 200 μL of the lysate was transferred to the next step, and excess lysate was stored in a clean 50ml tube in −20°C. To purify the DNA, 200 μL of the lysate was mixed with 400 μL in-house SPRI bead solution (Vesterinen et al. 2016) and purified using an Opentrons OT-2 automated liquid-handling robot (New York, USA). During the robotic steps, the DNA was bound to the SPRI beads, drawn to the magnet, and the supernatant was discarded. Then, the DNA pellet was washed twice using 40 μL freshly prepared 80% ethanol. After removing all ethanol, the pellet was dried, and DNA was eluted to 200 μL of pure RNAse, DNAse-free water. A DNA extraction control was added to each extraction batch, containing all the reagents, except the sample material. DNA purity and integrity were assessed by PCR success rate.

The fungal ITS gene region was amplified by using primer pair tagF-fITS7 (5’-GTGARTCATCGAATCTTTG-3’; Ihrmark et al. 2012) and tagR-ITS4 (5’-TCCTCCGCTTATTGATATGC-3’; White et al. 1990). This primer pair is designed to amplify fungi over plants (Høyer and Hodkinson 2021). All the primers included a linker-tag, enabling the subsequent attachment of NGS adapters. To increase the diversity of the amplicon library, each primer was used as two different versions, as including so-called heterogeneity spacers between the linker-tag and the actual locus-specific oligo. All PCR reactions were carried out as two technical replicates, and each replicate contained two heterogeneity versions of each primer. The reaction setup followed Kankaanpää et al. (2020) and included 5 μL of 2× MyTaq HS Red Mix (Bioline, UK), 2.4 μL of H_2_O, 150 nM of each primer (two forward and two reverse primer versions), and 2 μL of DNA extract per each sample in 10 μL total-volume. A blank PCR control was added to each PCR batch to measure the purity of reagents and the level of cross-contamination. PCR was performed under the following cycling conditions: 3 min in 95°C, then 35 cycles of 20 sec in 95°C, 30 sec in 55°C and 20 sec in 72°C, ending with 7 min in 72°C.

Library preparation followed Vesterinen et al. (2016) with minor modifications as follows: A dual indexing strategy was used, where each reaction (including technical replicates) was prepared with a unique combination of forward and reverse indices. All index sets were perfectly balanced so that each nucleotide position included either T/G or A/C, as this ensures base calling for each channel in the sequencing. For a reaction volume of 10 microliters, we mixed 5 μL of MyTaq HS RedMix, 500 nM of each tagged and indexed primer (i7 and i5) and 3 μL of the locus-specific PCR product from the first PCR. For library preparation PCR, the following protocol was used: initial denaturation for 3 min at 98°C, then 12 cycles of 20 seconds at 95°C, 15 seconds at 60°C and 30 seconds at 72°C, followed by 3 minutes at 72°C. All the indexed samples were then pooled and purified using magnetic beads (Vesterinen et al. 2018). Sequencing was done on an Illumina NovaSeq6000 SP Flowcell 2×250 (Illumina Inc., San Diego, California, USA) run, including PhiX control library by the Turku Centre for Biotechnology, Turku, Finland.

The bioinformatics pipeline closely followed Kaunisto et al. (2020). Paired-end reads were merged and trimmed for quality using 64-bit VSEARCH version 2.14.2 (Rognes et al. 2016). The primers were removed from the merged reads using software CUTADAPT version 2.7 (Martin, 2011), with a 20% rate for primer mismatches and 100 bp minimum length. The reads were then collapsed into unique sequences (singletons removed) with command ‘fastx_uniques’ using VSEARCH. Unique reads were denoised (i.e., chimeras were removed) and reads were clustered into ZOTUs (“ZOTU” = ‘(“ZOTU” = ‘zero-radius operational taxonomic unit) with command ‘unoise3’ using 32-bit USEARCH version 11 (Edgar, 2010). All samples with fewer than 50 reads in total were removed, and all ZOTUs from a sample with less than 20 reads for that ZOTU or with less than 0.05% of the total read number (all reads assigned to ZOTUs) of that sample were removed. Finally, ZOTUs were assigned to taxa by using the UNITE database (Abarenkov et al. 2020) with SINTAX (Edgar, 2010) in VSEARCH (Rognes et al. 2016). All ZOTUS with under 97% similarity to any reference database sequence were discarded. Finally, ZOTUS were assigned to a functional group by using the FUNGUILD python script (https://github.com/UMNFuN/FUNGuild), which matches taxonomic assignment against the FUNGuild database (Nguyen et al. 2016).

### Statistical analyses

#### Fungal communities of the soil vs. rhizosphere

To first characterize the level of specificity between fungi and plants (Figure 1: Box 1), we calculated a simple measure of interaction probability (Eq.1). To distinguish potential difference in specificity between functional guilds, this analysis was focused on the fungal genera that could be attributed to a specific functional group. In brief, we calculated the probability that an interaction occurs between a plant species *i* and a fungus *j* (i.e. that fungus *j* occurs on the rhizosphere of plant species *i*) at location *y (L_ijy_)*, given that fungus *j* is recorded within the soil around *i* at the same location *y (X_ijy_)*. For each plot, we divided the frequency with which each plant-fungus pair was found by the frequency of occurrence of the fungus within the soil:

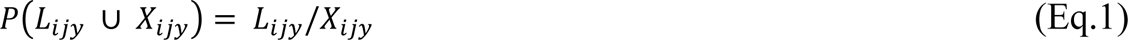

Rare fungi will easily be defined as specialists, since e.g. a fungus appearing only once on a plant will per necessity be observed on this plant species alone. Thus, we discarded fungal taxa occurring in less than 10% of the plots (i.e. in less than fourteen plots). However, many fungi were only detected within the rhizosphere while never being found within the soil (n=53, 10.6%), or were more often observed within the rhizosphere than in the soil, resulting in P(L_ijy_| X_ijy_)> 1 (n=57, 23.2%). As further experiments would be required to determine if this caused by obligate symbionts or molecular detectability issues, these fungi were discarded from the current analysis, as we were mainly interested in the recruitment of fungi from soil by the plants.

#### Biotic vs. abiotic effect on root associated fungi communities

In interpreting patterns of co-occurrence, it is important to note that P(L_ijy_| X_ijy_)=1 could arise either because fungus *j* is specialized on plant *i*, or because fungus *j* shares the exact same environmental preferences as the plant *i*. To disentangle these effects (Figure 1: Box 2), we used Hierarchical Modelling of Species Communities (HMSC, Ovaskainen et al. 2017, Ovaskainen and Abrego 2020). This model partitions the variation in species occurrences and co-occurrences to variation explained by the abiotic covariates vs. variation remaining after accounting for these abiotic effects, in the form of residual associations (Ovaskainen, Hottola, and Siitonen 2010; Ovaskainen et al. 2017). In this multivariate framework, a matrix of taxon-by-sample observations (the Y community-matrix, with entries y_ij_ for taxon *j* at plot *i*) is modelled as a function of a matrix of the plot-level environmental covariates (the X matrix, with entries x_ik_ for covariate *k* at plot *i*). As a basis for analyses, we created a data matrix of fungus taxa × plant taxa × plot × community type (soil or rhizosphere), resolved to the level of plant individuals. Following the terminology of Ovaskainen and Abrego (2020), this matrix is henceforth referred to as the **Y** community matrix. We note that any fungal genus *i* could potentially occur two times in the Y community-matrix, if recorded as present in both the soil and the rhizosphere. The two records were then treated as separate taxa and the community type as a trait (summarized in the T matrix, with entries “soil” for fungi recorded in the soil and “root” for fungi detected within the rhizosphere). Matrix Y was reduced to data on the presence/absence of taxa and fitted to a a probit HMSC model (Ovaskainen and Abrego 2020). Since very rare or very common species will contribute little information on the factors affecting species presence or absence, we included only genera that occurred in at least 40 samples (3%), in at least one of the datasets (soil or roots).

The occurrences of taxa were modelled as a function of edaphic conditions and bioclimatic variables relevant for fungal communities within the soil and within the rhizosphere (Burns et al. 2015; Zhang et al. 2016; Ni et al. 2018; Rasmussen, Bennett, and Tack 2020). As each sample corresponds to the fungal community found around a specific plant, or on its roots, we include plant identity as a categorical fixed effect, to test for differences in fungal incidence across plants. For each plot we included temperature (a continuous covariate), soil pH (a continuous covariate), soil Nitrogen content (%, a continuous covariate), and vegetation cover (%, a continuous covariate). To control for the effects of variation in sequencing depth, we included log(total number of reads) as a community type-specific covariate (with one value for the soil sample and one for the rhizosphere sample). All covariates were scaled to a mean of 0 and a variance of 1.

The HMSC models were fitted with the R-package Hmsc (Ovaskainen and Abrego 2020; Tikhonov et al. 2020). The models were fitted with eight chains in total, each with 1,500,000 iterations, which we discarded 500,000 as transient and which we thinned the remaining 1,000,000 by 4000 for a total of 2000 samples from the posterior. MCMC convergence was assessed by examining the potential scale reduction factors of the model parameters. The discriminatory power of the probit model was measured by calculating two different metrics, i.e. species-specific “areas under the curve” (AUC; Pearce and Ferrier 2000) and Tjur’s coefficient of discrimination (Tjur’s R^2^; Tjur 2012).

In the fitted models, the responses of taxa to fixed effects (as representing abiotic conditions) will inform us about the responses of fungal taxa to their environment, whereas the residual variance-covariance matrix will inform us about either biotic associations, or about joint responses to environmental covariates unmeasured in the study (Ovaskainen and Abrego 2020).

#### Network analyses

To quantify the relative impact of environmental conditions on partner selection in plant– fungus networks across the Arctic (Figure 1: Box 3), we quantified interaction structure using the H2’ index (Blüthgen, Menzel, and Blüthgen 2006) as a standardized measure of overall network specialization at the site scale.

##### Plant-fungus networks and partner selection across the Arctic

To quantify whether local (i.e., site-scale) networks show any clear ecological structuring pattern, we used a null model approach. To this aim, we generated randomly assembled networks based on random reallocation of links in the plant–rhizosphere association matrix of each local network using the “swap.web” algorithm of package bipartite (*v.2.18*; Dormann et al. 2018, Dormann 2023). This algorithm allows the removal of any systematic patterns while controlling the connectance (i.e maintaining the overall number of links in the network). Thus, we compared the observed network indices value (I_observed_) with the average value of the index across 1000 iterations of the null model (*I̅*_nulls_; Eq.2). We then expressed the difference between the null expectation and the observed value in units of standard deviations of the null distribution (σ_null_):

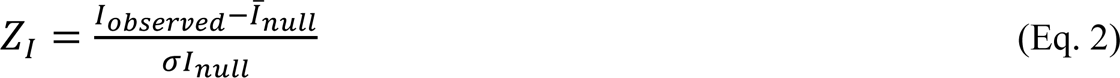

If the z score is higher than 2 or lower than −2, then we considered that the local network showed significantly more structure than expected by chance alone. Consequently, it would reflect a detectable imprint of ecological processes structuring plant–fungus interactions.

Null models can also be used to evaluate the role of functional traits in structuring ecological networks. To test whether the functional matching of interactions is stricter than expected under random associations, we applied the same approach as above to the subset of fungi, in each local network, to which functions could be assigned by matching with the FUNGuild database (Nguyen et al. 2016); see section *DNA-metabarcoding*). As an example, we extracted all the rhizosphere-associated fungi assigned to “mycorrhiza” and produced a local network containing only the interactions observed between these mycorrhizae and the different plants within a site (henceforth “the metaweb of functional mycorrhizae”). Again, we quantified the deviation (i.e z-score) between the value of the network index (i.e H2’) observed for plant–mycorrhiza and the mean expectation from randomized networks in units of standard deviations across 1000 iterations of the null model.

##### Does plant–fungus specialization change across environmental gradients?

To explore whether the level of specialization at the plot-level networks changed across environmental gradients (for a definition of plots vs. sites, see *Sample collection* section), we compared plot-level specialization to null models generated at the site level (and thus containing all the interactions observed within the *site*).

To generate the relevant null expectations under no association between the plot-level environment and the features of the local network, we randomized the full set of links in the plant-rhizosphere association matrix of each local network using the “swap.web” algorithm from the package bipartite (*v.2.18*; Dormann et al. 2018, Dormann 2023). Focusing on H2’, we then modelled the z-scores of the deviations between the H2’ values of the observed subweb to the null web as a function of bioclimatic variables (Estimated temperature, Nitrogen and Carbon content in the soil, pH and vegetation coverage). As above, we used the type of interactions (i.e the full network or the subnetworks consisting of mycorrhizal, saprotrophic or pathogenic fungi alone) as a categorical variable, to thereby test whether these follow similar patterns across environmental gradients.

For each site, we fitted a generalized linear mixed effects model (GLMM) with the measured environmental variables as fixed effects and the plot as a random effect – the latter to account for variation among plot-specific networks as due to e.g. differences in network size (i.e, number of plant species sampled, see *Sample collection*). These GLMMs were fitted using the package ‘glmmTMB’ in R (*v.1.1.7*; Brooks et al. 2017), using the following model specification:

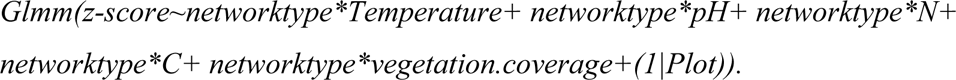

Each covariate was scaled and centred around 0. To validate model fit, we used functions implemented in package ‘DHARM’ (*v.0.4.6*; Hartig 2022). Model checks included visual inspection of residuals using quantile-quantile plots (to detect overall deviations from the expected distribution), with additional tests for adherence to the distribution assumed (KS test), for dispersion and for outliers. To identify systematic deviations from model assumptions, we also checked residuals plotted against predicted values.

## Results

### Fungal communities differ between the soil and the rhizosphere

The DNA sequencing produced a total of 384M reads reliably assigned to fungi. Out of this set, 156M sequence reads were retained for soil samples and 228M for rhizosphere samples. For soil samples, 7053 ZOTUs were reliably assigned to 444 genera of fungi; for rhizosphere samples, 7973 ZOTUs were reliably assigned to 459 genera (Appendix S1: Table S1). Rarefaction curves of fungi showed that the sequencing effort was sufficient among sites and community type (i.e. the sampling depth sufficed to recover more or less the full microbial communities within both the rhizosphere and the soil; Appendix S1: Figure S2). Out of the 459 fungal genera detected in rhizosphere samples, 278 were assigned to a functional group (60.6%). Among these fungal genera, 28 were assigned to ectomycorrhiza, one (genus *Oidiodendron*) to ericoid mycorrhiza, one (genus *Rhizophagus*) to arbuscular mycorrhiza, one (genus *Serendipita*) to orchid mycorrhiza and one (genus *Sebacina*) to a mycorrhizal type whose precise nature was uncertain. Additionally, 112 genera were assigned to saprotrophs and, 33 to plant pathogens.

At a Pan-Arctic scale, we found a similar diversity of fungal genera in the rhizosphere and in the soil (459 genera in rhizosphere vs. 444 in soil), with ratios ranging from 1.1:1 to 3.1:1 for fungal genera (Appendix S1: Figure S3). As a result, for each community type, a minority (13-41%) of the fungal genera detected was unique to the rhizosphere or to the soil (7-33%; Appendix S1: Figure S3). In contrast, a relatively large proportion of taxa (44-76%) was shared between the two communities (Appendix S1: Figure S3).

Multivariate ordination (NMDS) highlighted partial overlap, but also substantial differences between soil and rhizosphere fungal community composition (Figure 3). Fungal composition appeared more homogeneous within the rhizosphere than within bulk soil (Figure 3) – with little or no separation between plant species, but strong separation between presumed mycorrhizal type (Figure 3).

**Figure 3.**
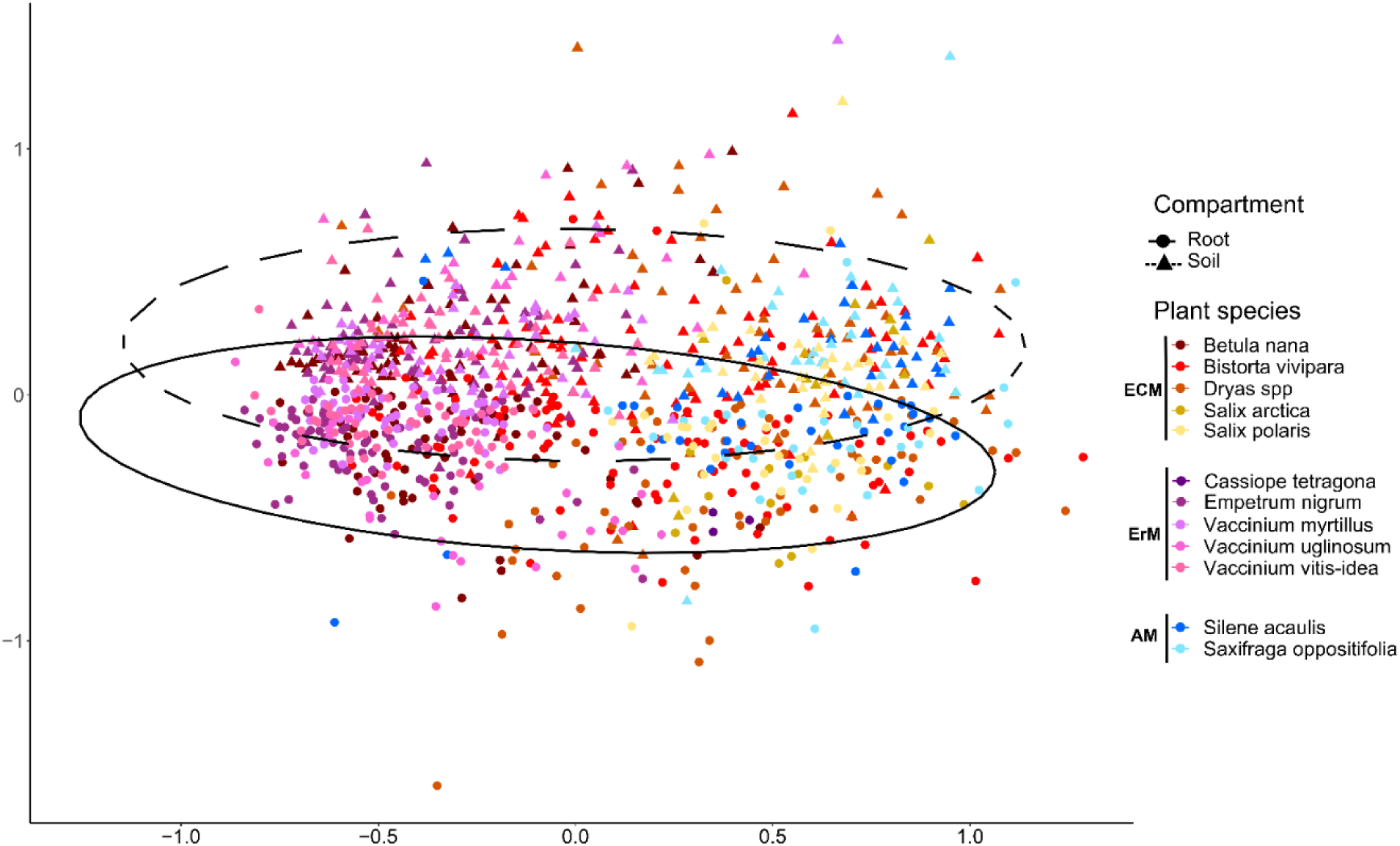
Multivariate ordination (NMDS) of fungal community composition found within the roots (filled line) of different plants species vs. bulk soil (dashed line). Ellipses summarize the within-community variability (75%) of each community type. Each color represents a plant species categorized into the type of mycorrhiza presumably associated with; ECM=Ectomycorrhiza, ErM=Ericoid mycorrhiza and AM=Arbuscular mycorrhiza. The NMDS was performed on species presence-absence data using the Jaccard distance. The stress value of the NMDS is 0.20; number of permutations = 1000, 2 dimensions, R^2^ = 0.82.

### Fungal recruitment varies across plants

In terms of the frequency of occurrence in the rhizosphere when present in the soil (Figure 1: Box 1), mycorrhizal fungi showed the highest incidence (Figure 4). These taxa were also characterized by the widest distribution across plant species, with several fungal genera showing high incidence across all plant taxa – regardless of the presumed mycorrhizal type of this plant. Endophytes and saprotrophic fungi formed an intermediary group, whereas pathogenic fungi were less frequently encountered in the rhizosphere and, were typically associated with only a few plant species.

**Figure 4.**
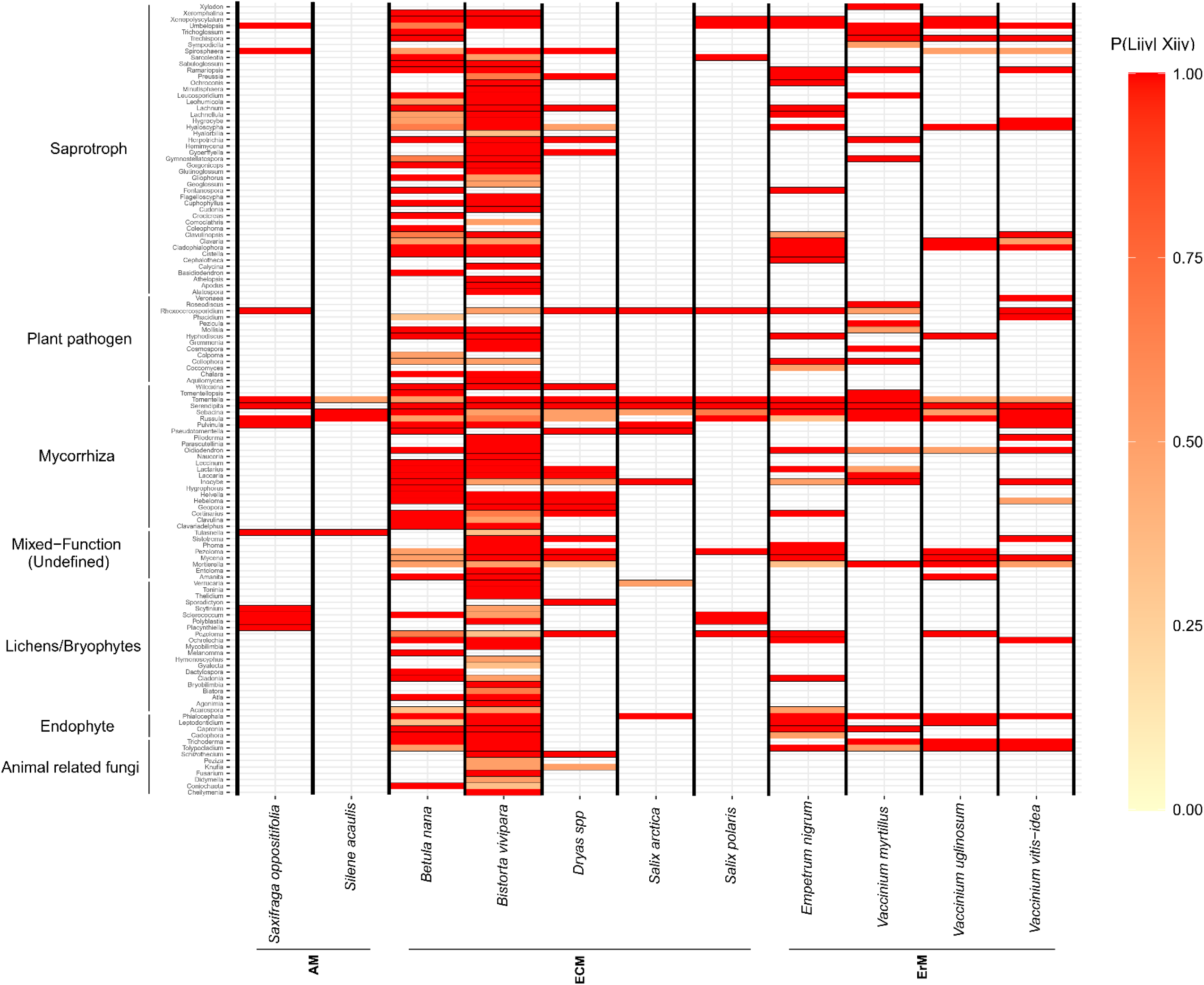
Frequency of association between fungal occurrence in the soil and in the rhizosphere around the same plants. As a simple measure of interaction probability, we show a heat map of the probability that an interaction between a plant species i and a fungus j occurs at location y (L_ijy_) when the fungus j is recorded within the soil around the same plant i at the same location y (X_ijy_). Each row represents a fungus assigned to a functional group and each column a plant species, with the color proportional to P(L_ijy_| X_ijy_). Plant species on the x axis are sorted according to their assumed mycorrhizal types, where AM= arbuscular mycorrhiza; ECM=ectomycorrhiza and ErM= Ericoid mycorrhiza.

Of the plants, *Bistorta vivipara* and *Betula nana* were the species with the highest proportion of strongly-associated fungi within their rhizosphere (Figure 4). Moreover, *Bistorta vivipara* showed frequent associations with almost every fungal genus (as assigned to a functional group) whenever the fungus was recorded within the soil (Figure 4). In contrast, *Silene acaulis* showed the lowest proportion of fungi strongly associated with their rhizosphere. For this species, only four fungal genera were frequently found in its rhizosphere when recorded in the soil (Figure 4).

### Fungi in the soil and in the rhizosphere respond similarly to their environment

The HMSC model used to quantify the relative importance of the abiotic and biotic environment in shaping the composition of fungi communities (Figure 1: Box 2) was successfully fitted to the data. MCMC convergence was satisfactory and the potential scale reduction factors were close to the theoretical optimum of one. The model achieved satisfactory discriminatory performance with a mean Tjur R^2^ of 0.22 and a mean AUC value of 0.90 (Appendix S1: TableS2). However, the explanatory power (Tjur R^2^) was similar amongst taxa present within the rhizosphere compared to the fungi present in the soil (rhizosphere: mean Tjur R^2^ = 0.22, AUC = 0.89; soil: mean Tjur R^2^ = 0.24, AUC = 0.90; Appendix S1: TableS2). Predictive performance assessed by cross-validation was lower than explanatory performance (mean cvTjur R^2^ = 0.16, mean cvAUC = 0.80). On average, these cross-validation indices were similar between community types (rhizosphere: mean cvTjur R^2^ = 0.13, cvAUC = 0.80; soil: mean cvTjur R^2^ = 0.14, cvAUC = 0.80s; Appendix S1: TableS2).

Overall, the bioclimatic environment explained the presence/absence of fungal taxa equally across community types (Appendix S1: TableS2). The random effects, especially at the sample level, explained the highest proportion of raw variance for both community types (rhizosphere=5.8%; soil=6.0%, Appendix S1: TableS2), followed by abiotic conditions (rhizosphere: 5.0%; soil=5.0%, Appendix S1: TableS2) and the host effect (rhizosphere: 2.9%; soil=2.2%; Appendix S1: TableS2). Similarly, across community types, pH accounted for the highest proportion of variation explained by abiotic conditions, followed by temperature (Appendix S1: TableS2). As expected, the host effect explained slightly more variance for the fungi within the rhizosphere than for fungi in the soil (Appendix S1: TableS2). Overall, the variance explained by individual covariates was largely consistent among different functional groups of fungi, with the strongest host effect for mycorrhizal fungi occurring within the rhizosphere (Appendix S1: TableS2)

In terms of responses to abiotic conditions, the occurrence of a taxon responded similarly to a given environmental feature across the two community types (soil vs rhizosphere; Figure 5) and we found no posterior support for a significant directional effect of the type of sample (i.e soil vs rhizosphere) on taxon-specific responses to environmental variables (Appendix S1: Figure S4-A). This was evidenced by consistency in the sign and significance of effects across the two communities (Figure 5; see Appendix S1: Figure S5, compare the color of juxtaposed tiles between community type within taxa).

**Figure 5.**
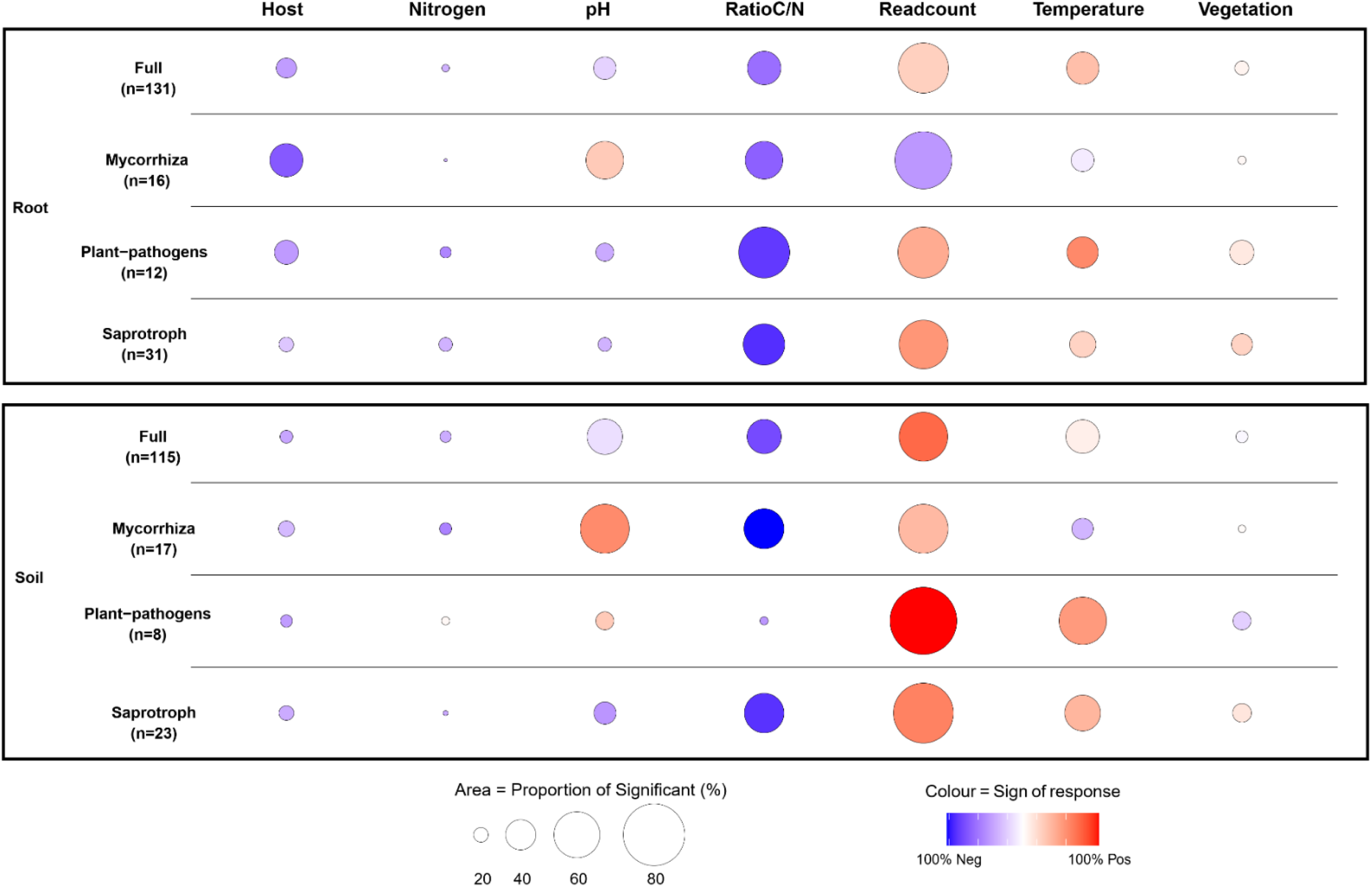
Summary of taxon-specific responses to environmental covariates for HMSC model of the presence-absence of fungal genera among community types. Circles size corresponds to the proportion of taxa for which a statistically supported response was detected (i.e. a response with at least 0.95 posterior probability), whereas circle colour shows the balance between negative and positive responses (see colour scale at bottom of graph; here white indicates that an equal number of taxa responded positively and negatively to the predictor in question. Statistical inference was based on the posterior distribution of the beta parameters, equivalent to regression coefficients. Ranges of values for individual covariates: Temperature: −11.9°C to +0.62°C; pH: 4.12–7.54; Nitrogen content (i.e., Nitrogen): 0.062-2.3**%**; Ratio Carbon/Nitrogen (i.e., Ratio C/N): 8.8-32.9**%**; Vegetation coverage (i.e., Vegetation): 5-99.4%; Sequencing depth, which is log transformed of total number of reads per sample (i.e., Readcount): 2.6-13.3).

Overall, only 31.3% of all fungal taxa detected within the rhizosphere and 19.9% of all fungi within the soil responded significantly to the identity of the host plant. By comparison, 32.1% of the fungi detected in the rhizosphere and 32.5% of taxa detected in the soil responded significantly to abiotic covariates (i.e Temperature, Nitrogen content, pH, vegetation coverage and the C:N ratio; Figure 5). Among these abiotic covariates, the Carbon/Nitrogen ratio elicited the strongest response, with 40.8% and 54.1% of fungal taxa in the soil and rhizosphere, respectively, showing significant and mainly negative responses (Figure 5). The next most frequent response was observed for temperature, to which 46.1% of fungal taxa in the soil and 37.9% of fungi in the rhizosphere showed a significant and predominantly positive response (Figure 5). To pH, fungi within the rhizosphere and the soil again showed similar responses, with mixed positive and negative responses among individual taxa (Figure 5). Overall, both the variance-partitioning results (Appendix S1: TableS2) and strong heterogeneity in the estimated beta parameters revealed taxon-specific responses to the environmental covariates (Figure 5; Appendix S1: Figure S5). In a similar vein, community-level predictions showed consistency in predicted species richness along environmental gradients (Appendix S1: Figure S6). Across host plant species, the predicted richness of fungi in the rhizosphere was higher than richness predicted in the soil (Appendix S1: Figure S6-A). Across environmental gradients, the predicted richness of fungal taxa responded similarly between community types (Appendix S1: Figure S6B-F). In terms of residual covariance, we found little evidence for biotic interactions (Appendix S1: Figure S4-B). Thus, fungal occurrence in the soil and in the rhizosphere seems governed by abiotic imprints.

### Plant-rhizosphere networks are relatively non-specialized across the Arctic and mainly driven by local environmental conditions

Fungal interaction partners were widely shared between plants (Figure 1: Box 3; Figure 6A). Across plants, roughly one-third of the fungi found within the rhizosphere were unique to a specific plant even when plants co-occurred within the same plot (Figure 6B). The overall proportion of fungi shared among co-occurring plants was relatively evenly distributed among plant pairs (Fig 6B). However, the proportion of shared fungi showed a clustering between presumed mycorrhizal type of the plant species, i.e. arbuscular and ericoid mycorrhizal plants shared only few fungi, while ectomycorrhizal plants shared as many fungi with plants presumed to be associated with arbuscular mycorrhiza as with plants presumed to be associated ericoid mycorrhiza (Figure 6B). The highest proportion of fungi shared in the rhizosphere was found between *Cassiope tetragona* and *Empetrum nigrum* (21.2%; Figure 6B). *Empetrum nigrum* shared a particularly high number of partners with *Vaccinium vitis-idea*, *Betula nana* and *Vaccinium myrtillus* (Figure 6A). *Salix arctica* and *Vaccinium uglinosum* showed the highest rates of unique non-shared fungi, with 48.4% and 45.7% of all fungi uniquely found within their rhizosphere (Figure 6B). In terms of functional groups of fungi, the majority of fungi unique to a plant were assigned to saprotrophs, whereas the majority of fungi shared between plants were assigned to symbiotic or antagonistic functions (i.e mycorrhiza, endophytes and pathogenic fungi; Appendix S1: Figure S7). Mycorrhizal fungi represented around a third of fungi commonly shared between plant species, whereas pathogens accounted for around 15% of shared fungi (Appendix S1: Figure S7).

**Figure 6.**
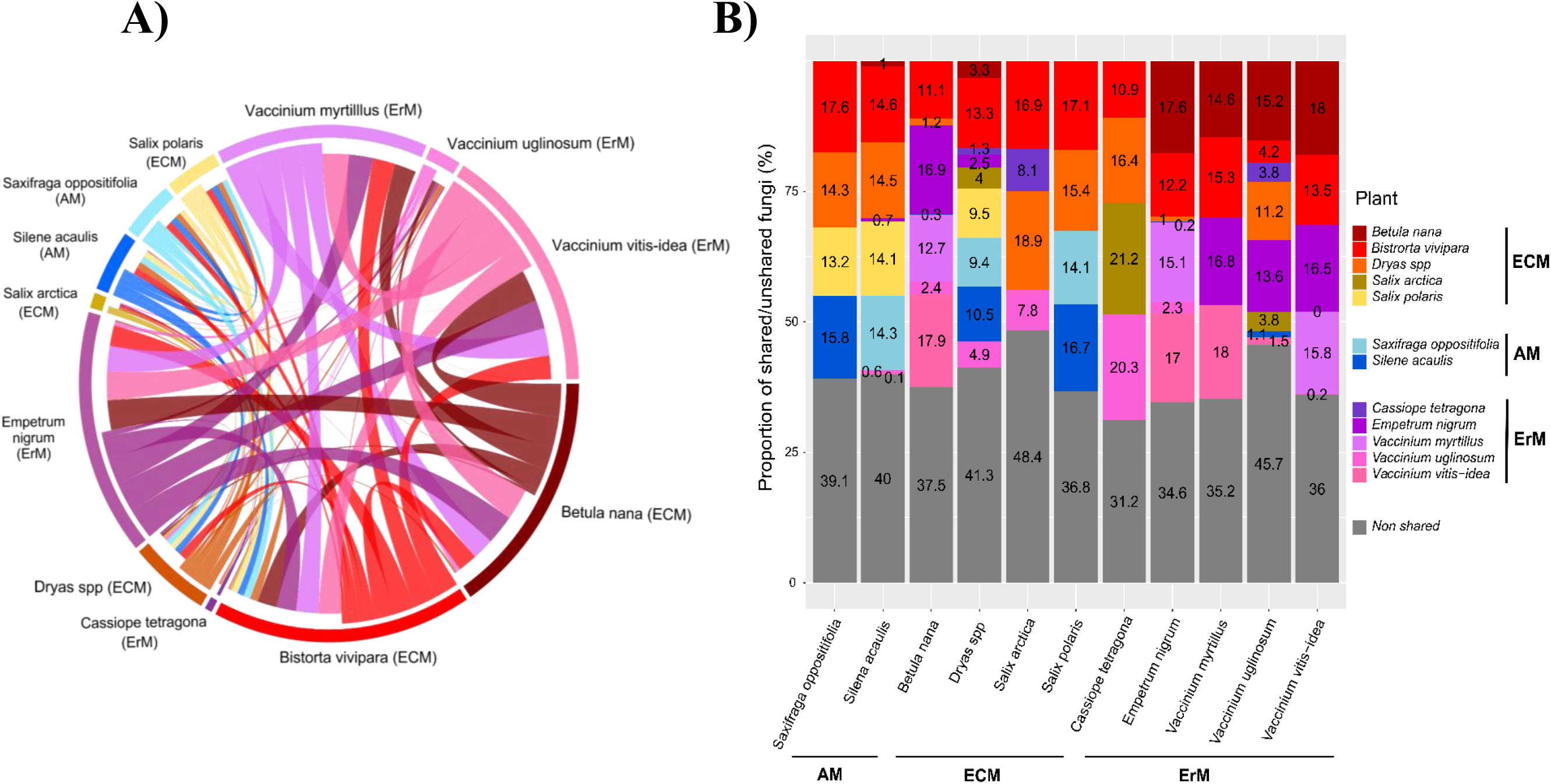
Overlap in rhizosphere-associated fungi between co-occurring plants. For each plot across all our sites, we quantified the number of rhizosphere-associated fungi recorded for each combination of plant species. Panels A) and B) show the numbers and proportions of rhizosphere-associated fungi shared between two plant species when present in the same plot. Panel (A) is a chord diagram showing overlap between specific plant pairs, whereas B) decomposes the number of all rhizosphere-associated fungi detected on a plant species (individual columns) into the specific proportion unique to this plant (in grey) versus shared with other plant species (colored sections). Plants species in panel B) are sorted according to their assumed mycorrhizal types where AM=arbuscular mycorrhiza; ECM=ectomycorrhiza and ErM= Ericoid mycorrhiza.

To further quantify if the local host preferences for each site varies across the Arctic, we explored the site-level network specialization network index (as quantified by H2’). The H2’ index varied considerably between ecological groups (i.e between the mycorrhiza and plant-pathogens) and between sites. The deviation between observed values and values expected under the null model slightly increased with latitude, but few networks appeared significantly more structured than expected by random (Figure 7A). The mean structure of sub-networks including only mycorrhiza or plant pathogens did not differ from random expectations (Figure 7A). Although specialization was not significantly different from what we can expect by chance, the patterns shifted over space: with increasing latitude, deviations from the null model became increasingly negative for the mycorrhizal network, while they became increasingly positive for the plant-pathogen networks. In other words, mycorrhizal and saprotrophic networks tended to become more generalized with increasing latitude, while the pathogenic network and the full networks tend to become more specialized than expected by chance. Thus, we further investigated the level of specialization across local environmental gradients (Figure 7B). Standardized estimates from the GLMMs (Figure 7B) identified soil pH and temperature as the strongest predictors of local specialization, with a strong site-specificity (Figure 7B). Moreover, the responses of the different network compartments (i.e the full network vs. plant–mycorrhizal and plant–pathogenic interactions) differed strongly from each other in terms of their responses to environmental conditions (Figure 7B).

**Figure 7.**
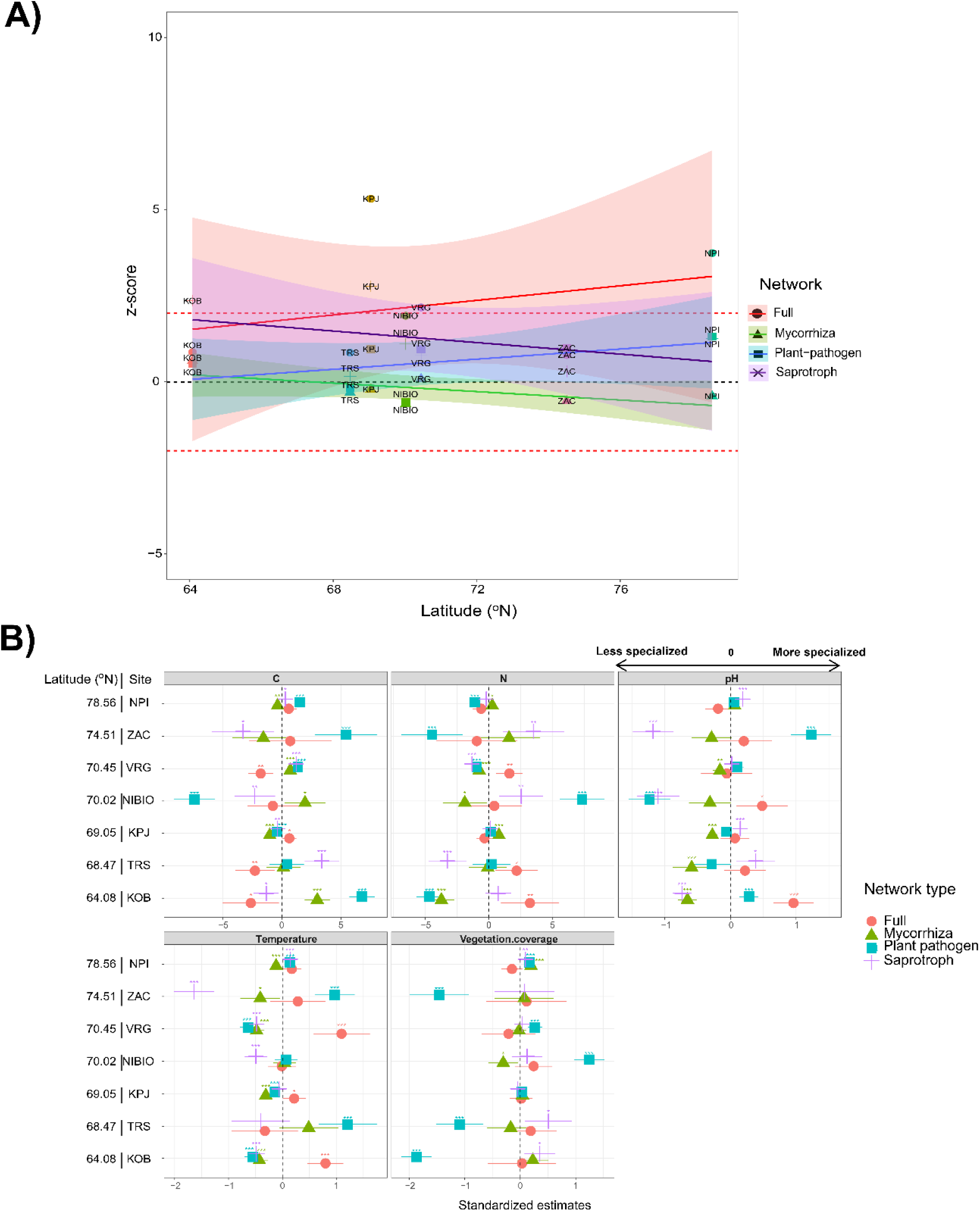
Deviations from randomness in the specialization of networks as a function of latitude (Panel A) and across local environmental gradients (Panel B). Panel A) shows the z-score, i.e. the deviation (in units of standard deviations) between the observed H2’specialization index and the expected value, as derived from 1000 realizations of a null model. If the z-score is higher than 2 or lower than −2, then the observed specialization is significantly different from the value expected under the null model. Panel B) shows dot-and-whisker plots of the responses of network specialization to different features of the environment across sites. Shown are standardized estimates of site-specific glmm models of specialization (measured by z-scores between plot specialization and the randomized specialization of the network at site level) as functions of variation in Carbon, Nitrogen, pH, temperature and Vegetation coverage within each site. Each color and shape represents a standardized estimate of a specific type of networks, with estimates ordered by latitude along the y axis. In both panel A) and B) “Full network” represents the full plant-fungus matrix, whereas “Mycorrhiza” represents the site-specific sub-network including only fungi assigned to the functional group of mycorrhiza. “Plant-pathogens” represent the site-specific sub-network including only fungi assigned to the functional group of plant-pathogens. Site acronyms: KPJ= Kilpisjärvi, Finland; VRG= Varanger Peninsula, Norway, NIBIO= Gandvik valley, Norway; KOB= Kobbelfjord, South-West Greenland; TRS= Toolik Research Station, USA; NPI= Norsk Polarinstitut, Ny Alesund, Svalbard; ZAC= Zackenberg, North-East Greenland.

## Discussion

Given the importance of associations between plants and soil fungi, the two should be studied in conjunction. In this study, we dissect the effects of biotic and abiotic drivers on networks consisting of plants and rhizosphere associated fungi across the Arctic. While the fungal composition of the soil and the rhizosphere proved significantly different from each other, the occurrence of a fungus in the plant rhizosphere was strongly dependent on its presence in the soil. Within both the rhizosphere and the soil, fungal occurrence was mainly driven by abiotic environmental conditions, with little effect of host plant or biotic interactions. In terms of the interaction networks of fungi and plants, we found a generally low but variable level of host specificity of fungi. Variation in specialization across the Arctic was mainly driven by conditions at the local rather than the regional scale. At the local scale, temperature, pH and vegetation coverage appeared to be the main environmental factors impacting network specialization. With increasing pH and temperature, the networks tended to be more generalized in nature. Below, we will discuss each of these findings in turn.

### Soil- and root associated fungal communities respond similarly to their environment

As our first key objective (Figure 1: Box 1), we assessed how the probability of plant– fungus interaction is modulated by the occurrences of the fungi within the soil and by the bioclimatic environment, respectively. Based on global patterns (Soudzilovskaia et al. 2015; Bahram et al. 2018; Tedersoo et al. 2022), we expected congruent changes in the composition of soil and rhizosphere-associated fungal communities. As an alternative hypothesis, we proposed differential change in the two community types (i.e rhizosphere and soil). If plants provide a strong “physical buffer” (Sikes 2010), then we would expect differences in how the fungal communities of the rhizosphere vs. fungi present in the soil respond to environmental gradients. More explicitly, soil fungi would be mainly driven by abiotic conditions, while rhizosphere-associated fungi would be mainly driven by biotic environment.

Of these alternatives, the first expectation was clearly borne out by our results, whereas the second was refuted. Overall, the fungal composition of the rhizosphere was significantly different from that of the soil, and the rhizosphere of plants sustained a higher diversity of fungi than did the soil. We found 54 fungal genera unique to roots (11% of the full set of fungi) and 34 fungal genera unique to soil (8% of the full set of fungi). However, the results of our joint species distribution (HMSC) model revealed that fungal occurrence in both communities was mainly driven by pH and temperature (Appendix S1: Table S2). When coded as a trait, community types as such (soil vs. rhizosphere) did not modify species responses. Neither did we detect any signs of strong residual associations among rhizosphere-associated fungi, after accounting for the abiotic effects (Appendix S1: Figure S9-A), suggesting a general lack of pronounced biotic interactions among fungi. Along environmental gradients, predictions of emergent community features (here: overall species richness) were largely consistent among community types.

In the current results, there are little if any suggestions of biotic interactions among fungal taxa as being strong drivers of community assembly. Only two mycorrhizal fungi from the genus *Geospora* and *Lactarius* showed a positive residual association between community types (i.e., rhizosphere vs soil), suggesting that the occurrence of these genera in the soil is positively associated with their presence within the roots in a way not accounted for by their environment (Appendix S1: Figure S5-B). This partly contrasts with previous studies suggesting more pronounced associations between fungal guilds (Hannula and Träger 2020). In sites characterized by low fertility (which is generally the case in Arctic ecosystems), abundances of mutualists (mycorrhizal fungi) and saprotrophs have been found to be negatively related to each other, suggesting antagonistic competition for nutrients (Hannula and Träger 2020). In our case, the residual co-occurrences (i.e., after considering the effect of environment) of fungi in the rhizosphere showed a sparse pattern (Appendix S1: FigS9-B). Only 1% of all possible associations between saprotrophic and mycorrhizal genera were detectably positive and 1% negative.

Overall, these findings attest to a lack of strong biotic imprints on either community types, and rather suggests that abiotic drivers (i.e., environmental filtering) dominate fungal community assembly across both the arctic soil and the rhizosphere. Our current results are thus well in line with previous reports suggesting that fungal richness in the rhizosphere is more closely linked to abiotic than biotic variables (Blaalid et al. 2014; Alzarhani et al. 2019).

### Root communities are weakly impacted by plant identity

As our second key objective, we aimed to evaluate fungal partner selection among plant species (Figure 1: Box 2). Here, we found little imprint of host identity, but rather a strong signal of opportunistic partner selection among taxa present in the soil. Thus, arctic fungi appear to show largely opportunistic patterns of associations with individual plants. In support of this inference, the presence of individual fungal taxa in the soil had a strong influence on their presence in the plant rhizosphere, and individual plant species shared massive amounts of fungi with each other. Low levels of host specificity extended to all functional groups (i.e. to fungal taxa identified as saprotrophs, mycorrhiza, plant pathogens and endophytes; Appendix S1: Figure S8).

The current findings may appear surprising, given that we had *a priori* selected plants representing different types of fungal associations (ectomycorrhiza, arbuscular mycorrhiza, ericoid mycorrhiza etc.). Nonetheless, individual fungi showed relatively low fidelity to these groups. Overall, out of all 459 root-associated fungal genera here detected across the Arctic, only 15% were unique to a plant species and 85% were found within the rhizosphere of at least two different plant species. Among plant mycorrhizal types, we found no clustering of shared fungi between plant species that are known to be associated with similar mycorrhizal types (Figure 3; Figure 6). In fact, the distribution and proportion of shared fungi between plant species was relatively even (Figure 6B-C). However, we observed two distinct clusters in root associated fungi between plant presumed to be associated with arbuscular and ericoid mycorrhiza (Figure 3; Figure 6A-B).

Among the plants selected for this study, *Silene acaulis* is assumed to associate with arbuscular mycorrhiza (Abrego et al. 2020; Rasmussen et al. 2022). In the current study, we found relatively few arbuscular mycorrhiza fungi in the rhizosphere of this species – whereas the fungal genera that we did detect were also found in plants known to form ectomycorrhizal associations, such as *Salix polaris* (Botnen et al. 2014, Arraiano-Castilho et al. 2020; Figure 4). While ectomycorrhizal fungi are frequently detected on plants originally defined as lacking mycorrhiza or being associated with arbuscular mycorrhiza (Kytöviita 2005), we were surprised to systematically record *Oidiodendron* in the rhizosphere of *Bistorta vivipara* (cf. Figure 4). *Bistorta vivipara* is assumed to be mainly colonized by ectomycorrhizal fungi or sometimes by arbuscular mycorrhiza (Eriksen, Bjureke, and Dhillion 2002), whereas *Oidiodendron* belongs to the group of ericoid mycorrhiza. This finding underscores the complexity of plant-fungus interactions and suggests that the associations between plants and mycorrhizal fungi may be more nuanced and variable than previously thought.

Overall, the opportunistic sharing of fungi among plants detected is in line with several studies from arctic and alpine areas, which have shown that the dominant plants associated with ectomycorrhiza or ericoid mycorrhiza share the same pool of fungal symbionts (Timling et al. 2012; S. Botnen et al. 2014; S. S. Botnen et al. 2020; Toju and Sato 2018). To frequently associate with different plants occurring in the same plot can reflect efficient host foraging behavior (Lekberg et al. 2010). Indeed, such opportunistic foraging can be an efficient strategy to find alternative hosts and/or increase bargaining power in harsh environments (Lekberg, Hammer, and Olsson 2010; Chagnon, Bradley, and Klironomos 2020). From a plant perspective, *Betula nana* and *Bistorta vivipara* showed frequent associations with almost every fungal genus assigned to a functional group. This may reflect a high degree of generalism on the plants’ part, or even opportunistic behavior. Highly generalist plants have been considered as key regulators of ecosystem dynamics (e.g., promoters of community “stability” or “robustness”; (Chagnon 2016; Chagnon, Bradley, and Klironomos 2020)). As an example, *Betula nana* may potentially facilitate the revegetation by shrubs and possibly the expansion of trees into the Arctic by maintaining ectomycorrhiza communities during tundra fire (Hewitt et al. 2013; Timling et al. 2014). Surprisingly though, we found *Silene acaulis* – a plant species mainly recorded in arctic, harsh environments – to be the plant species associated with the lowest proportion of the fungal taxa of the surrounding soil, suggesting high selectivity of partners. This finding runs opposite to our initial expectations, since in a harsh environment, association with any available partner tends to be better than being associated with no partner at all (i.e., as there is a high cost to partner rejection (Steidinger and Bever 2014; Chagnon, Bradley, and Klironomos 2020).

In conclusion then, our study paints a clear picture of arctic plants and fungi as being generalist in terms of partner preferences. However, whether or not specific taxa will co-occur and interact is mainly driven by environmental conditions.

### Plant-fungus networks are mainly structured by local environmental gradients

Until recently, networks involving plants and microorganisms belowground have received little attention – and to this date, plant–fungus networks of the Arctic remain poorly known (Bahram, Harend, and Tedersoo 2014; Wong et al. 2023). As our third key objective, we aimed to describe and explore the processes that shape plant-fungi networks in the Arctic (Figure 1: Box 3). At a regional scale (i.e., at the site level; Figure 2), the overall preference of fungi for plant interaction partners was no higher than expected by chance. Neither at the level of the full root-associated community nor at the level of functional groups did we find consistent evidence for host specificity. This lack of specificity is in line with previous studies reporting low specialization among root-associated fungi and plants in the Arctic ( Botnen et al. 2014; Monard et al. 2016). The lack of network specialization and of partner preferences suggest that environmental conditions might override the effects of the host plant in fungal community assembly (Timling et al. 2014; Maciá-Vicente and Popa 2022; Wong et al. 2023)

In this context, we found that the deviation of the local networks (i.e the plot-level associations) from the randomized site-level metaweb was strongly structured by environmental gradients (Figure 7B). This shift into generalist networks points to a re-arrangement of local networks under environmental change. Overall, our findings offer support for the “stress gradient hypothesis” where, under increasingly harsh environmental conditions, resources are primarily allocated to survival and reproduction rather than to competition – avoiding the risks associated with host specialization (Botnen et al. 2014; Tylianakis and Morris 2017)

Interestingly, partial networks consisting of mycorrhizal, pathogenic and saprotrophic associations alone showed lower network-level interaction specificity than did full networks encompassing plants and all fungi. This finding resonates with the results of Toju et al. (2018) from temperate Japanese forests. There they reported higher specialization at the level of the full interaction network than at the level of the partial networks of mycorrhizal and saprotrophic associations. Nonetheless, our analyses of different compartments of the networks were clearly challenged by the difficulties in assigning fungal taxa to functions. As mentioned above, 40% of fungal genera could not be assigned to any functional group. Thus, the pooling of these different unidentified interaction types may generate different conclusions regarding the structure of the full plant-fungi networks. Clearly, much more work is needed to improve references database and define the ecological role of arctic fungi before we can fully comprehend ecological networks in these ecosystems.

## Conclusions

Our analyses reveal complex relationships between the environment and fungal community structure and diversity. Local communities in the plant rhizosphere are formed from the species pool available in the local soil, with strong imprints of the environment on the species present. Nonetheless, among locally co-occurring plant and fungal species, we found major variation in plant–fungus associations and low levels of partner specificity. This level of plasticity suggests high adaptability of plant–fungus networks in the face of environmental changes. As our current study is clearly observational in nature, we urge experimental studies exploring the rewiring achieved under controlled conditions.

## Author contributions

BP, NMS and TR designed the study. BP, MT, PEA and TR collected the biological samples. BP measured a part of the edaphic variables used in this study. Bioname, led by EV, generated metabarcoding libraries from root and soil samples and produced fungal community data. BP, AC, EVG and TR designed the statistical analyses. BP and JS analyzed the data, with input from TR, AC, HW, NMS, EVG. BP, TR and HW wrote the first draft of the manuscript. All authors then helped revise the manuscript or provided comments on the final manuscript.

## Conflict of Interest

The authors declare no conflict of interest.

## Funding

B.P and T.R were funded by the Academy of Finland (VEGA, grant 322266 to T.R). TR was also funded by the European Research Council (ERC) under the European Union’s Horizon 2020 research and innovation programme (ERC-synergy grant 856506— LIFEPLAN). BP was further supported by a grant from the Ella and Georg Ehrnrooth foundation. Field work was supported by an INTERACT Transnational and Remote Access grant (Call 2020, project VEGA). AC was founded by Kone Foundation. J.S. was supported by the Academy of Finland’s ‘Thriving Nature’ research profiling action

## Acknowledgments

We are indebted to all collaborators and field assistants from the Norsk Polarinstitut; Toolik Research Station (Amanda Young); NIBIO-Svanhovd research station station (Juho Vuolteenaho and Helena Klöckener), as well as Minna Viljamaa for participating in the generation of empirical samples. We are grateful to Marjo Kilpinen and Eija Takala for their help in measuring pH. We wish to acknowledge DNA analysis company Bioname for their expertise and help regarding the molecular analysis and interpretation of the results. We acknowledge CSC-IT Center for Science Ltd., Espoo, Finland, for the allocation of computational resources. We thank Pablo de la Peña Aguilera, Antoine Becker-Scarpitta and Giovanni Strona for stimulating discussions and early input on this study. Laura Antaõ, Maria Pires-Braga, Jussi Mäkinen, Sonja Saine, and Bess Hardwick offered important input in improving an earlier draft of the manuscript.

---

**Table S1.**
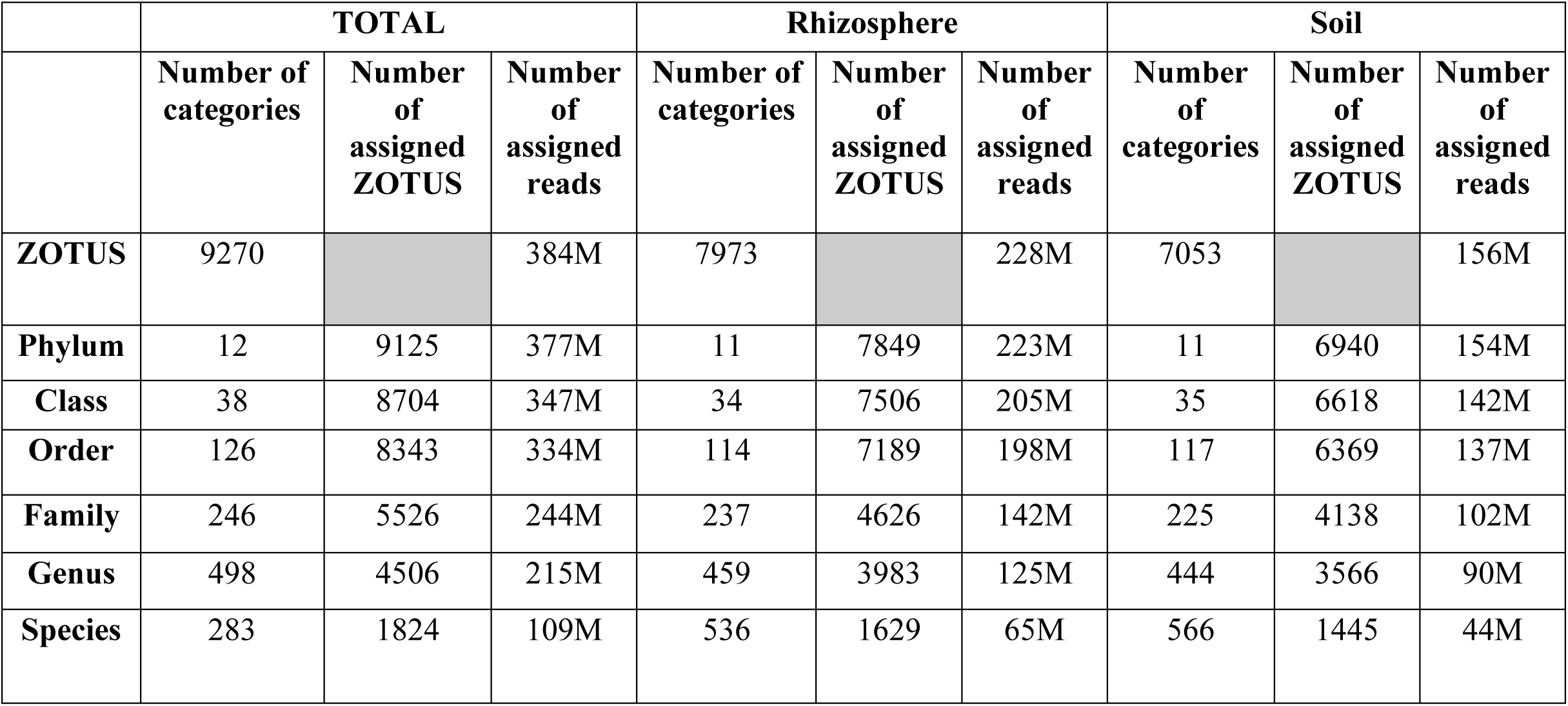
Summary of taxonomic and functional assignment success for each compartment (i.e., root vs soil) Each entry identifies the number of sequences reliably assigned to a taxon at the respective taxonomic level for the compartment (rhizosphere or soil) in question.

|  | TOTAL |  |  | Rhizosphere |  |  | Soil |  |  |
| --- | --- | --- | --- | --- | --- | --- | --- | --- | --- |
|  | Number of categories | Number of assigned ZOTUS | Number of assigned reads | Number of categories | Number of assigned ZOTUS | Number of assigned reads | Number of categories | Number of assigned ZOTUS | Number of assigned reads |
| **ZOTUS** | 9270 |  | 384M | 7973 |  | 228M | 7053 |  | 156M |
| **Phylum** | 12 | 9125 | 377M | 11 | 7849 | 223M | 11 | 6940 | 154M |
| **Class** | 38 | 8704 | 347M | 34 | 7506 | 205M | 35 | 6618 | 142M |
| **Order** | 126 | 8343 | 334M | 114 | 7189 | 198M | 117 | 6369 | 137M |
| **Family** | 246 | 5526 | 244M | 237 | 4626 | 142M | 225 | 4138 | 102M |
| **Genus** | 498 | 4506 | 215M | 459 | 3983 | 125M | 444 | 3566 | 90M |
| **Species** | 283 | 1824 | 109M | 536 | 1629 | 65M | 566 | 1445 | 44M |

**Figure S1.**
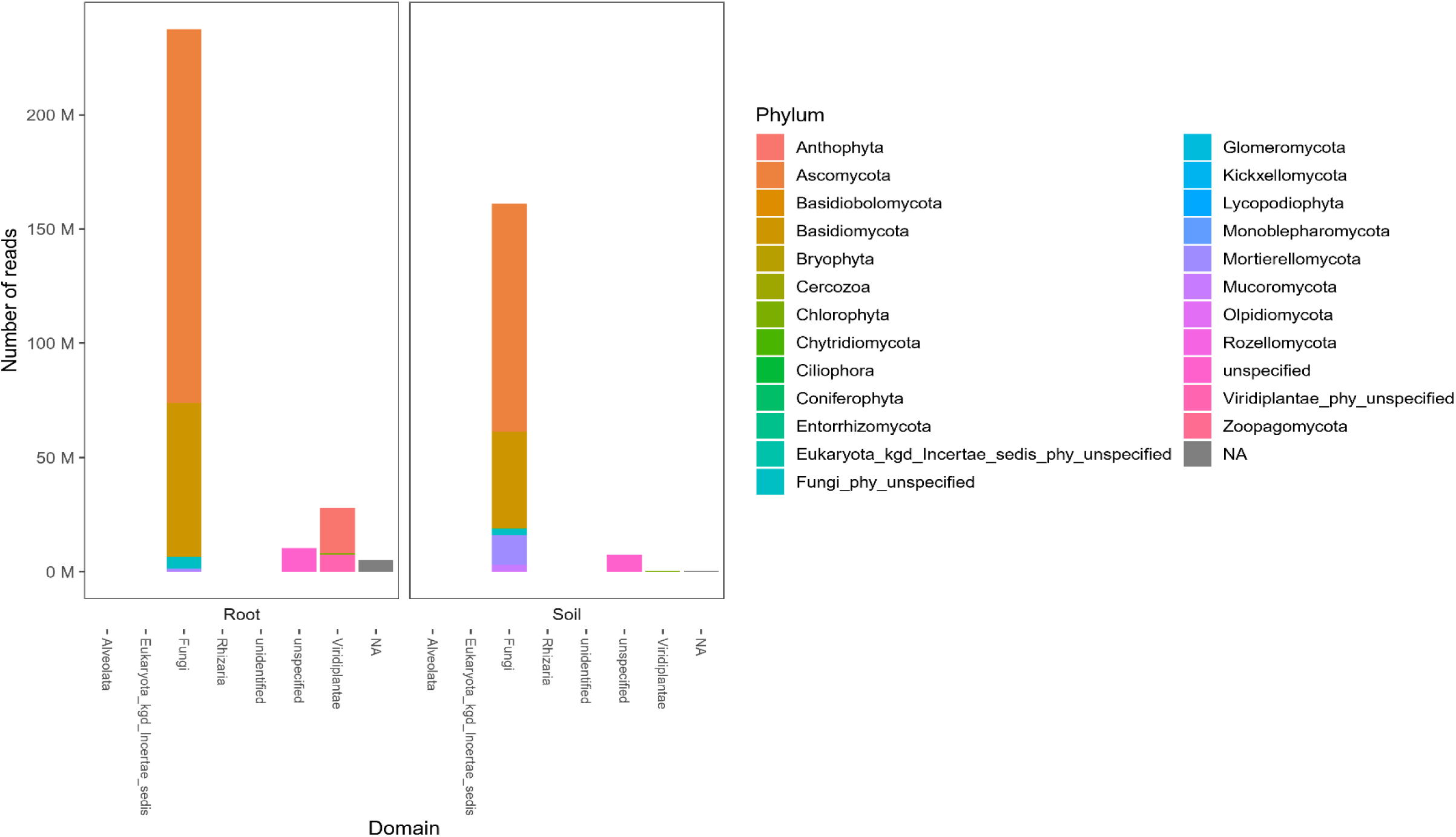
Proportion of reads assigned to fungi vs. other eukaryote domain. The x axis shows the major domains of Eukaryote. Bar colours reflect the composition of phyla detected. Reads assigned to Plantae or Unspecified were discarded from the downstream analyses.

**Figure S2.**
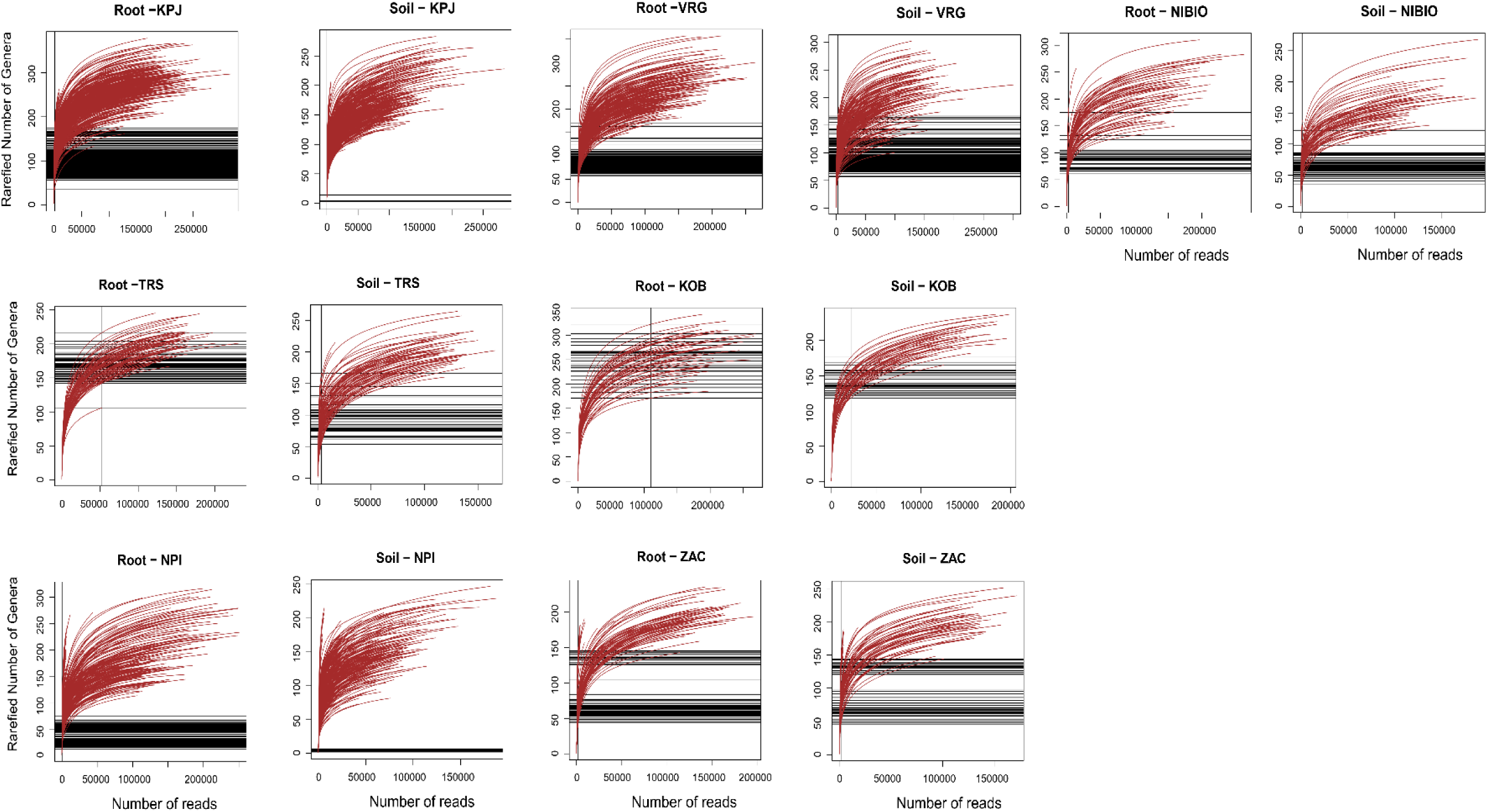
Rarefaction curves of the number of fungal genera per sample as a function of read number. Sites: KPJ= Kilpisjärvi, Finland; VRG= Varanger Peninsula, Norway, NIBIO= Gandvik valley, Norway; KOB= Kobbefjord, South-West Greenland; TRS= Toolik Research Station, USA; NPI= Ny Alesund, Svalbard; ZAC= Zackenberg, North-East Greenland. The curves shown were drawn with the vegan package (Oksanen, 2010) in R.

**Figure S3.**
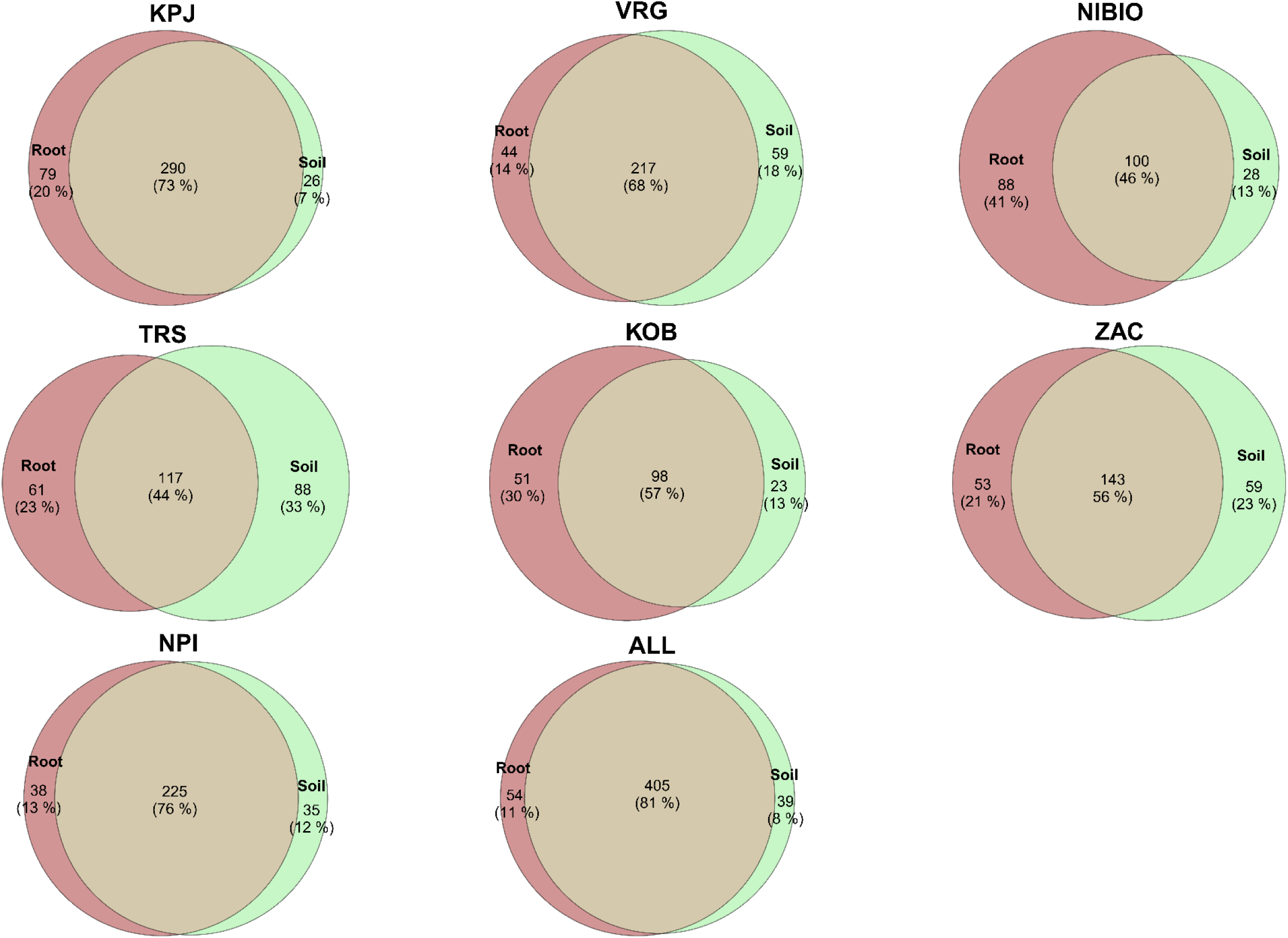
Euler diagrams showing the shared and unique fungal genera detected in root or soil samples. Numbers within the intersections identify the number of fungi detected in both compartments. KPJ= Kilpisjärvi, Finland; VRG= Varanger Peninsula, Norway, NIBIO= Gandvik valley, Norway; KOB= Kobbefjord, South-West Greenland; TRS= Toolik Research Station, USA; NPI= Ny Alesund, Svalbard; ZAC= Zackenberg, North-East Greenland. All= All sites pooled together

**Table S2.** Discrimination success achieved for the HMSC model of our fungal communities, for each compartment (soil vs. rhizosphere) The left-hand part of the Table summarizes the discriminatory power of the model, as based on two indices: Tjur R^2^ and AUC (see main text for definitions). Two aspects are evaluated : explanatory power (as reflected by Tjur R^2^ and AUC) and predictive power (as reflected by cross-validation Tjur R^2^ and cross-validation AUC). The right-hand part of the Table shows the average proportion of variation explained by each variable included in the model.

|  | Fungal functional group | Model performance indices |  |  |  | Predictors and raw variance explained |  |  |  |  |  |  |  |  |  |
| --- | --- | --- | --- | --- | --- | --- | --- | --- | --- | --- | --- | --- | --- | --- | --- |
|  |  | Tjur R<sup>2</sup> | AUC | Cross-validation Tjur R<sup>2</sup> | Cross-validation AUC | Temperature | Soil pH | Nitrogen content | Ratio C/N | Vegetation coverage | Readcount | Host | Random: Site | Random: Plot | Random: Sample |
| **Full model** | **Total (n=246)** | 0.22 | 0.9 | 0.16 | 0.8 | 1.2 | 2.1 | 0.2 | 0.7 | 0.5 | 3.4 | 2.9 | 3.1 | 2.4 | 5.8 |
| **SOIL** | Total (n=115) | 0.24 | 0.9 | 0.14 | 0.8 | 1.1 | 2.3 | 0.3 | 0.8 | 0.5 | 4.1 | 2.2 | 3.9 | 2.6 | 6 |
|  | Mycorrhiza (n=17) | 0.26 | 0.89 | 0.15 | 0.78 | 1.3 | 2.7 | 0.3 | 1.2 | 0.5 | 2.0 | 3.4 | 3.6 | 2.4 | 9.1 |
|  | Plant pathogens (n=8) | 0.16 | 0.88 | 0.08 | 0.77 | 0.8 | 1.4 | 0.2 | 0.4 | 0.6 | 2.6 | 2.3 | 1.5 | 2.2 | 4.0 |
|  | Saprotroph (n=23) | 0.24 | 0.89 | 0.15 | 0.79 | 1.2 | 2.4 | 0.3 | 1.0 | 0.7 | 4.4 | 2.6 | 2.8 | 3.1 | 5.7 |
| **ROOT** | Total (n=131) | 0.22 | 0.89 | 0.13 | 0.8 | 1.3 | 2.1 | 0.3 | 0.7 | 0.6 | 2.8 | 2.9 | 2.6 | 2.3 | 5.8 |
|  | Mycorrhiza (n=16) | 0.21 | 0.90 | 0.12 | 0.79 | 1.3 | 2.6 | 0.2 | 0.8 | 0.7 | 2.1 | 5.6 | 2.0 | 2.3 | 10.5 |
|  | Plant pathogens (n=12) | 0.22 | 0.89 | 0.14 | 0.81 | 1.5 | 1.4 | 0.2 | 0.7 | 0.5 | 3.3 | 2.7 | 1.7 | 2.9 | 4.8 |
|  | Saprotroph (n=31) | 0.24 | 0.89 | 0.15 | 0.8 | 1.1 | 2.7 | 0.4 | 0.9 | 0.9 | 2.7 | 3.3 | 2.5 | 2.5 | 5.4 |

**Figure S4.**
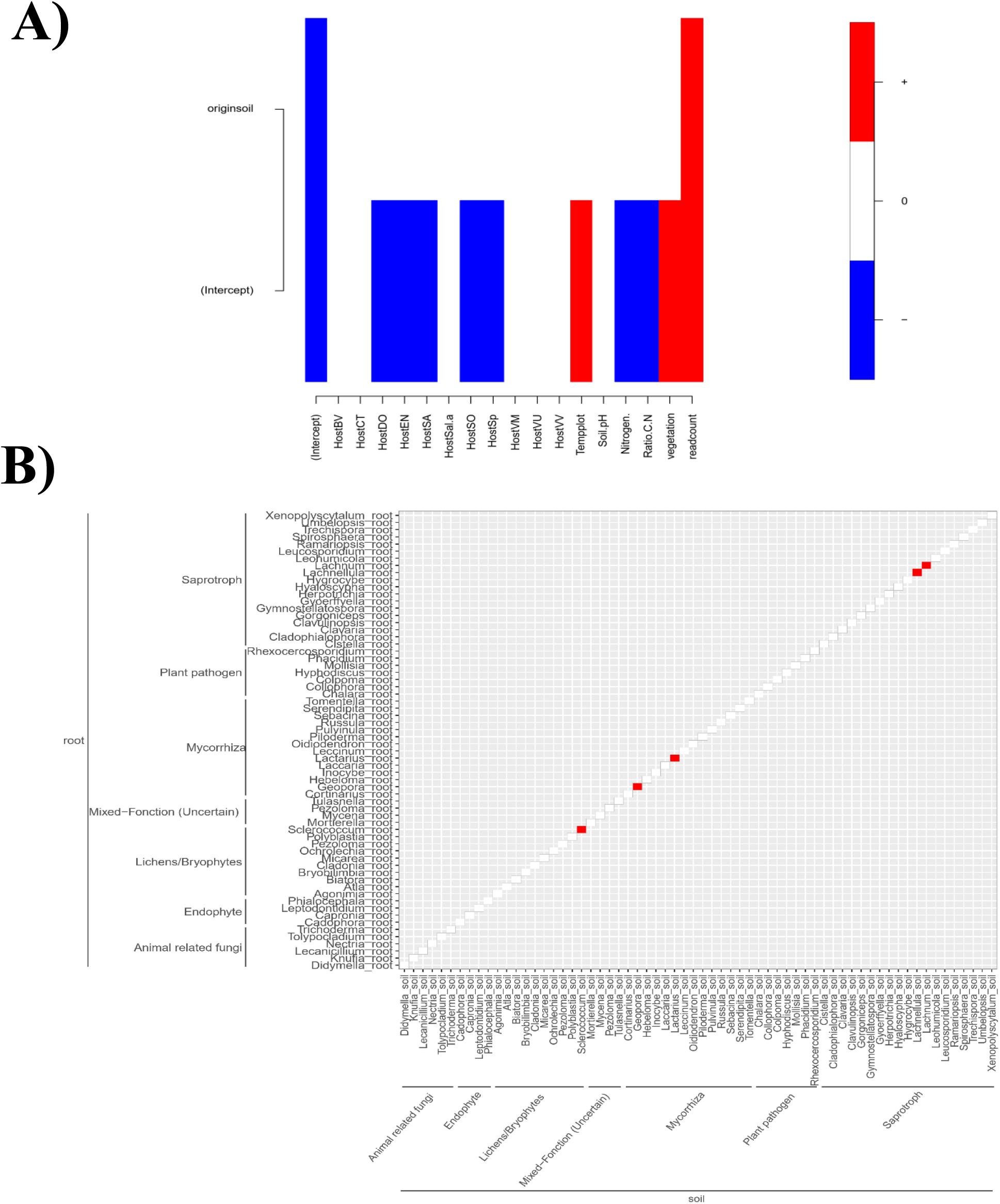
Estimates of A) consistency in response among fungi in soil and rhizosphere and B) biotic imprints. The top panel (A) shows estimates of the gamma parameter, as reflecting the impact of the trait (i.e. soil vs rhizosphere) on the estimated response. The intercept corresponds to rhizosphere associated fungi. Tile colours indicate whether a group of organisms is responding stronger (red) or weaker (blue) to a specific covariate, when compared to rhizosphere-associated fungi (posterior support >0.95). The bottom panel (B) represents the estimated pairwise residual associations among fungal taxa scored in both compartments (i.e., in the soil and in the rhizosphere). Here, significant residual association will suggest that the occurrence of the species in the soil and in the rhizosphere is more strongly associated (red) or dissociated (blue) stronger than accounted for by the environmental covariates measured. (Note that all entries fall on the diagonal, as each fungal taxon is shown as a line and a column

**Figure S5.**
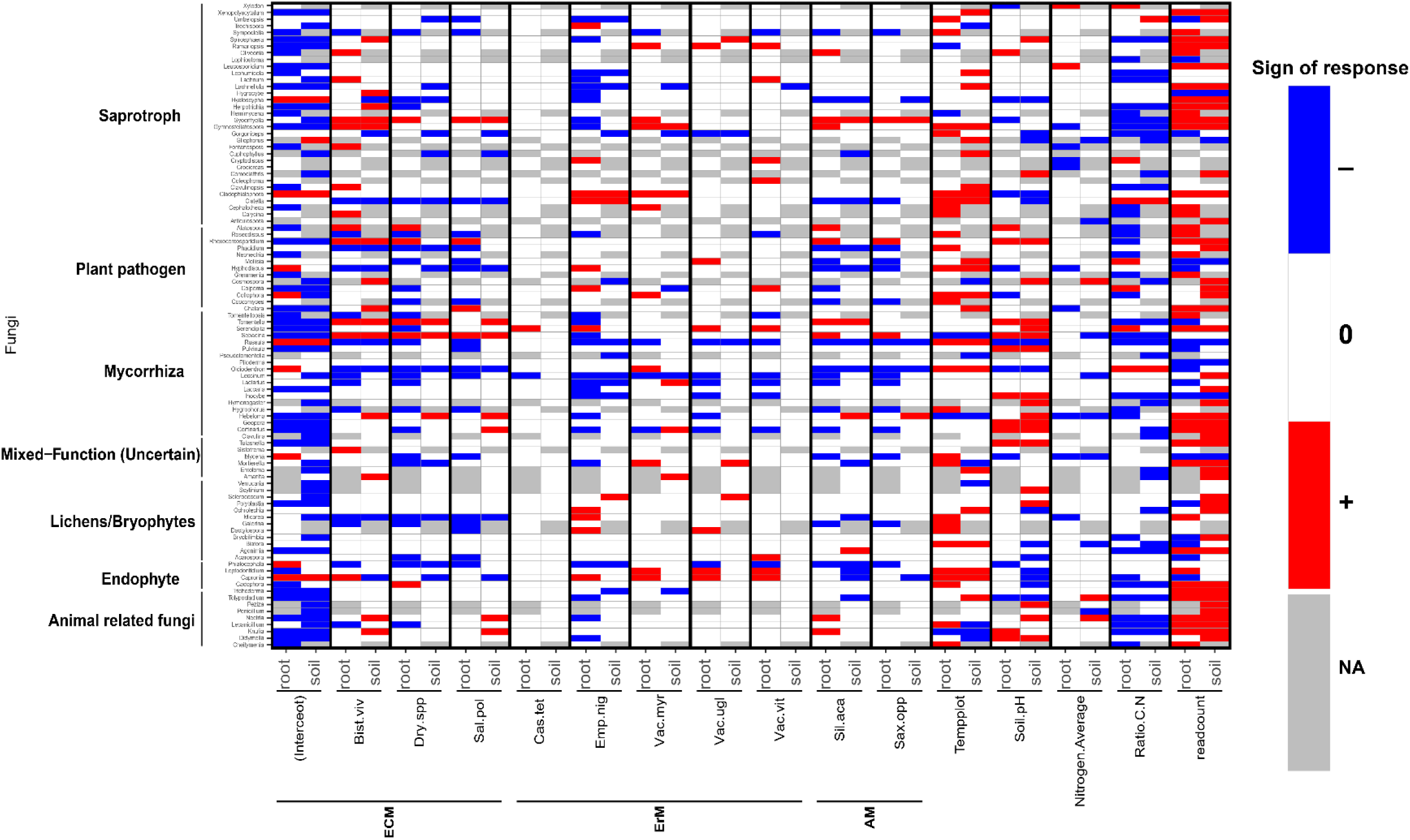
Beta parameter estimates (i.e., taxon-specific estimates of environmental responses, equivalent to regression coefficients) for HMSC models of the presence-absence of fungal genera among compartments. Each row indicates a fungal genus assigned to a functional group. Each tile represents the response of an individual taxon to a given driver: a host plant (columns 1-12), an abiotic condition (columns 13-16) or the sample-specific read count (column readcount). The intercept is the baseline of a specific fungus, i.e. in an average site in plant *Betula nana*, with covariates set to their averages and the values of random effects at set to. Juxtaposed tile-pairs within black horizontal dividers separate the response of the fungus in the rhizosphere (left-hand tile) vs. the soil (right-hand tile). Grey tiles indicates that a fungus has not been recorded in the associated compartment. Blue and red colours indicate parameter values that were estimated to be negative and positive, respectively, with at least 0.95 posterior probability. Fungi are sorted according to the functional group assigned to them (see labels along the y-axis). Plants species in the x axis (columns 1-12) are sorted according to their assumed mycorrhizal types whereas AM= arbuscular mycorrhiza; ECM=ectomycorrhiza and ErM= Ericoid mycorrhiza. HostBV=*Bistorta vivipara*; HostDO=*Dryas* spp; HostSp=*Salix polaris*; HostCT=*Cassiope tetragona*; HostEN=*Empetrum nigrum*; HostVM=*Vaccinium myrtillus*; HostVU=*Vaccinium uglinosum*; HostVV=*Vaccinium vitis-idea*; HostSA=*Silene acaulis*; HostSO=*Saxifraga oppositifolia*.

**Figure S6.**
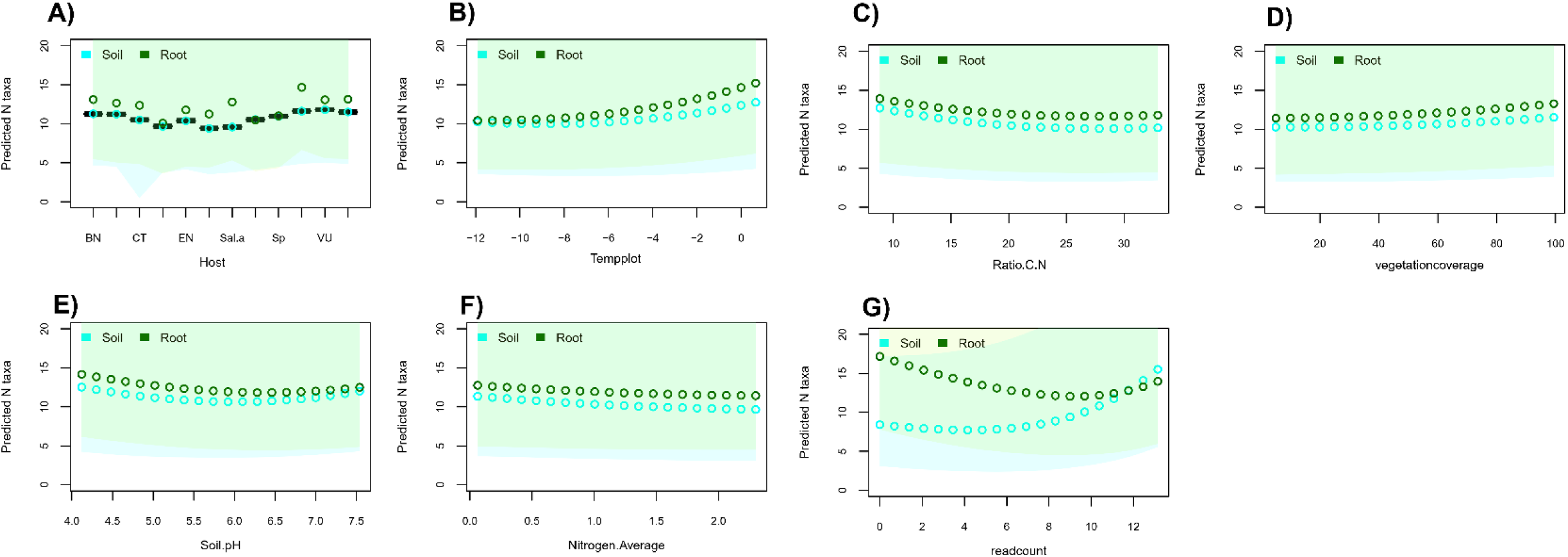
Posterior predictions of the mean numbers of fungal genera across compartments (soil in green vs. rhizosphere in cyan) in relation to each driver studied. (A) Host, where each category represent an unique plant species; (B) Temperature; (C) C:N-ratio; (D) Vegetation coverage; (E) pH; (F) Nitrogen content; and (G) sequencing depth (i.e., read count).

**Figure S7.**
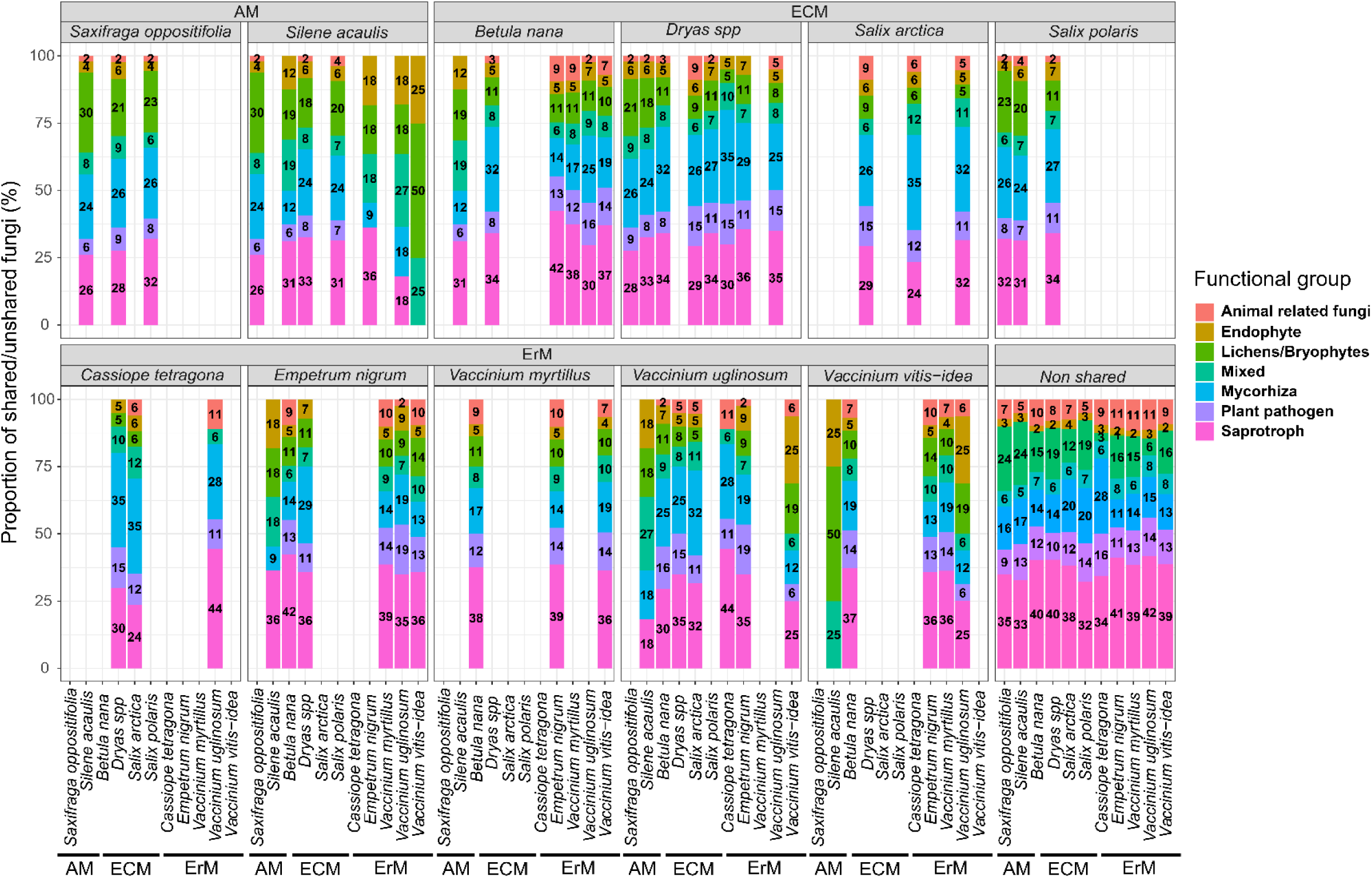
Composition of functional groups shared between plant species co-occurring in the same plot. Each panel corresponds to a specific plant species, each bar shows the whole set of fungi shared between two plants, and each colour distinguished the proportion of shared fungi assigned to a particular functional group. Plants species are sorted according to their assumed mycorrhizal types whereas AM= arbuscular mycorrhiza; ECM=ectomycorrhiza and ErM= Ericoid mycorrhiza.

## Text S1. Quantification of plant partner preferences

To examine whether network properties can be attributed to the ecological requirement of the individual plants and their associated fungi (i.e partner selectivity; Vaughan et al. 2018), we compared the preferences of each fungal genus observed in the rhizosphere to expectations from null models generated with the ‘generate_null_net’ function from the econullnetr package (Vaughan et al. 2018). More specifically, we used the default function, which uses interaction frequency as a proxy of interaction strength. Then, the ‘generate_null_net’ function generates null models following the algorithm of (Agustí et al. 2003), which is based upon interaction data and independent estimates of plant availability (i.e plants collected per each site) whereas the total number of links for each networks (i.e degree) is held constant (see Vaughan et al., 2018 for more details). These null models are then compared against the observed interactions between plant and fungi (to ascertain the extent to which resource choice deviated from random (i.e. density dependence). In this context, we compared the frequencies of observed to expected links (with the latter produced by null models for each site) for each pair of fungal genus and plant species. To identify whether our different plant species are more or less often associated with a specific functional group of fungi, we used the standardised effect size (SES) values obtained from the econullnetr package (Fig.S8).

---

**Figure S8.**
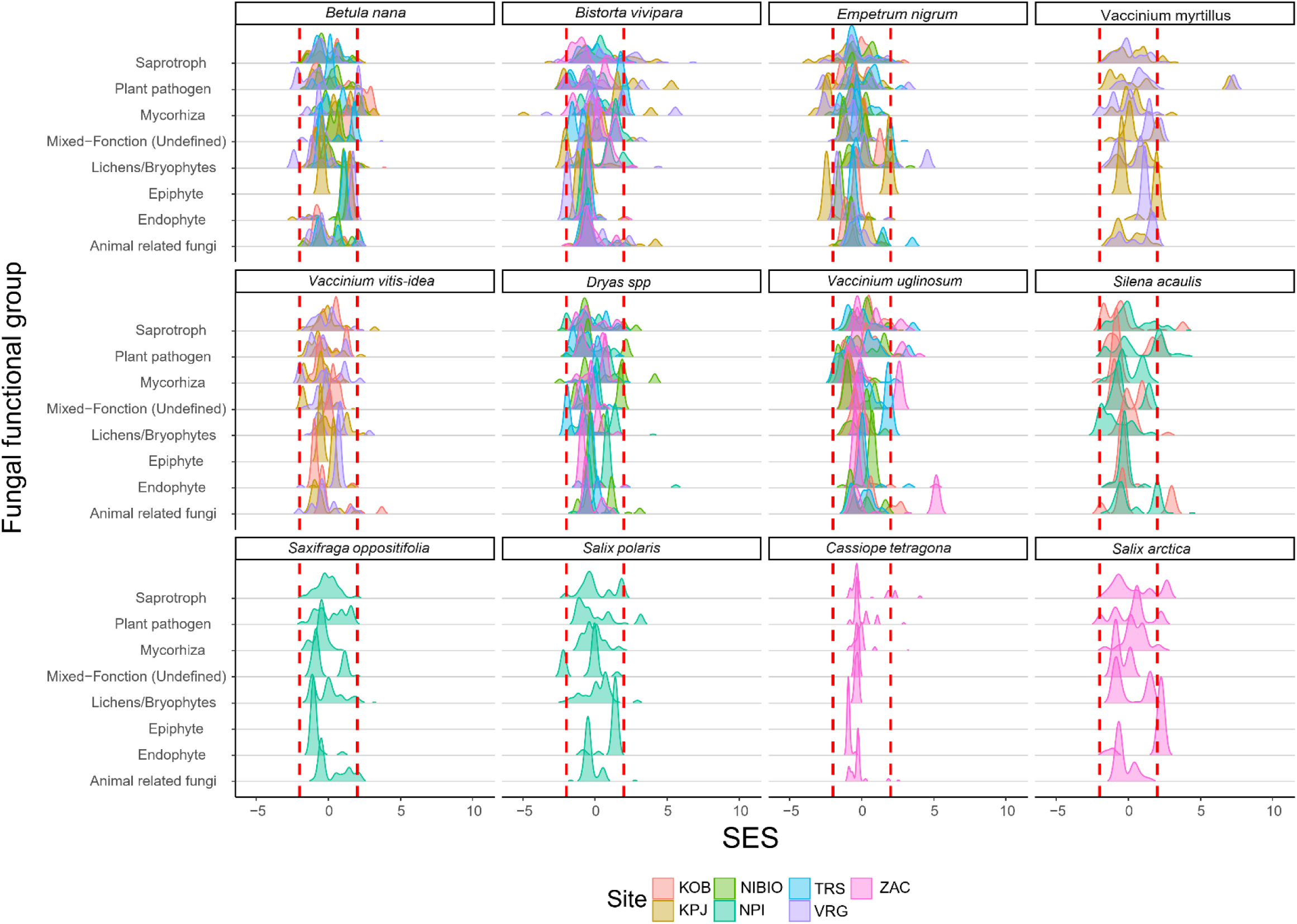
The preferences of different functional groups of rhizosphere-associated fungi quantified by SES. Shown are deviations between the observed number of plant-fungus links (aggregated by functional group) compared to frequencies expected under the null model. Deviations larger than 2 indicate that the observed number of links between a fungus and the plant is significantly higher than the number of links expected by chance. The density plots represent the distribution of all fungi aggregated by functional group. Site-specific acronyms: KPJ= Kilpisjärvi, Finland; VRG= Varanger Peninsula, Norway, NIBIO= Gandvik valley, Norway; KOB= Kobbefjord, South-West Greenland; TRS= Toolik Research Station, USA; NPI= Ny Alesund, Svalbard; ZAC= Zackenberg, North-East Greenland-For details, see Text S1.

**Figure S9.**
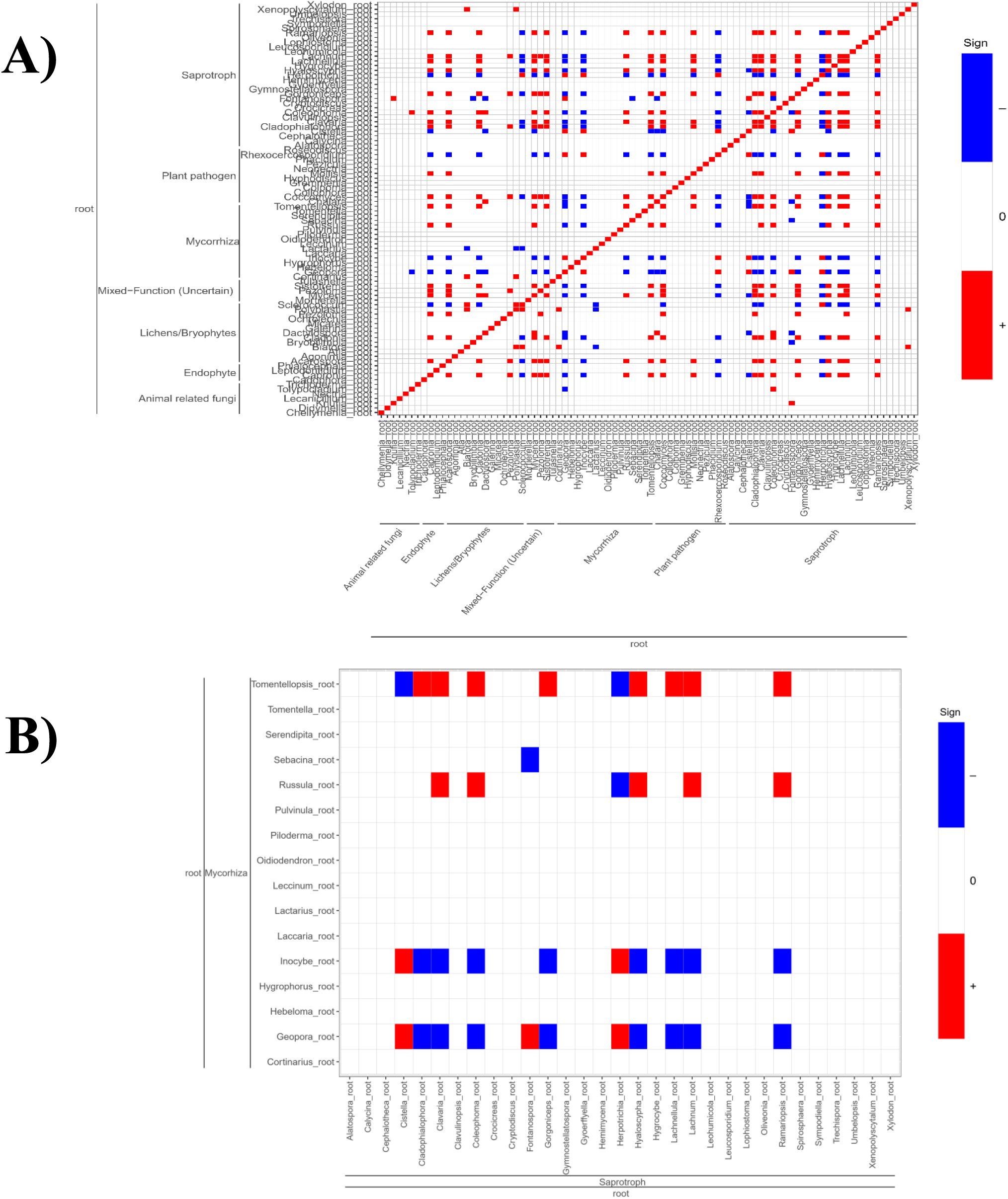
Residual associations detected amongst different functional groups of fungi detected in the roots. The panel A) shows estimated pairwise residual associations amongst fungal genera found within the root only. Fungi are ordered by functional groups with taxa sorted in the same order in both axis. Filled cells indicate species pairs showing association with at least 90% posterior probability, with blue for negative associations, red for positive associations.The panel B) shows the rresidual associations detected among saprotroph and mycorrhizal fungi found within the rhizosphere. Rows correspond to individual saprotrophic fungal genera in the rhizosphere, and columns to the mycorrhizal genera found within the rhizosphere. Filled cells indicate species pairs showing association with at least 95% posterior probability, with blue for negative associations, red for positive associations.

## Notes

### Competing Interest Statement

The authors have declared no competing interest.

